# AKAP150-anchored PKA regulation of synaptic transmission and plasticity, neuronal excitability and CRF neuromodulation in the lateral habenula

**DOI:** 10.1101/2023.12.06.570160

**Authors:** S.C. Simmons, W.J. Flerlage, L.D. Langlois, R.D. Shepard, C. Bouslog, E.H. Thomas, K.M. Gouty, J.L. Sanderson, S. Gouty, B.M. Cox, M.L. Dell’Acqua, F.S. Nugent

## Abstract

Numerous studies of hippocampal synaptic function in learning and memory have established the functional significance of the scaffolding A-kinase anchoring protein 150 (AKAP150) in kinase and phosphatase regulation of synaptic receptor and ion channel trafficking/function and hence synaptic transmission/plasticity, and neuronal excitability. Emerging evidence also suggests that AKAP150 signaling may play a critical role in brain’s processing of rewarding/aversive experiences. Here we focused on an unexplored role of AKAP150 in the lateral habenula (LHb), a diencephalic brain region that integrates and relays negative reward signals from forebrain striatal and limbic structures to midbrain monoaminergic centers. LHb aberrant activity (specifically hyperactivity) is also linked to depression. Using whole cell patch clamp recordings in LHb of male wildtype (WT) and ΔPKA knockin mice (with deficiency in AKAP-anchoring of PKA), we found that the genetic disruption of PKA anchoring to AKAP150 significantly reduced AMPA receptor (AMPAR)-mediated glutamatergic transmission and prevented the induction of presynaptic endocannabinoid (eCB)-mediated long-term depression (LTD) in LHb neurons. Moreover, ΔPKA mutation potentiated GABA_A_ receptor (GABA_A_R)-mediated inhibitory transmission postsynaptically while increasing LHb intrinsic neuronal excitability through suppression of medium afterhyperpolarizations (mAHPs). Given that LHb is a highly stress-responsive brain region, we further tested the effects of corticotropin releasing factor (CRF) stress neuromodulator on synaptic transmission and intrinsic excitability of LHb neurons in WT and ΔPKA mice. As in our earlier study in rat LHb, CRF significantly suppressed GABAergic transmission onto LHb neurons and increased intrinsic excitability by diminishing small-conductance potassium (SK) channel-mediated mAHPs. ΔPKA mutation-induced suppression of mAHPs also blunted the synaptic and neuroexcitatory actions of CRF in mouse LHb. Altogether, our data suggest that AKAP150 complex signaling plays a critical role in regulation of AMPAR and GABA_A_R synaptic strength, glutamatergic plasticity and CRF neuromodulation possibly through AMPAR and potassium channel trafficking and eCB signaling within the LHb.

## 1. Introduction

The scaffold protein A-kinase anchoring protein 79/150 (79 human/150 rodent/Akap5 gene) is a crucial regulator of synaptic receptor trafficking, synaptic transmission and plasticity, and neuronal excitability by anchoring protein kinases (e.g., protein kinase A, PKA, and protein kinase C, PKC) and phosphatases (e.g., calcineurin, CaN) and other signaling molecules (e.g., the transcription factor nuclear factor of activated T-cells, NFAT) to subcellular nanodomains at specific synapses (glutamatergic and GABAergic) ^1–9^ and ion channels [e.g., M-type potassium channels, A-type potassium channels, transient receptor potential vanilloid 1 (TRPV1) and L-type calcium channels] ^10–16^. AKAP150 anchoring of PKA, PKC and CaN has been shown to mediate the opposing effects of these enzymes in postsynaptic trafficking of both AMPARs and GABA_A_Rs during glutamatergic and GABAergic plasticity ^1–9^. For example, phosphorylation of GluA1 subunit of AMPARs with AKAP150-anchored PKA is required for stabilization and insertion of AMPARs in the synapse and promotion of long-term potentiation (LTP), while AKAP-anchored CaN dephosphorylates GluA1 subunit of AMPARs and removes AMPARs from the synapse which is required for long-term depression (LTD) in hippocampal CA1 neurons^17^. Similarly, transient recruitment of GluA2-lacking calcium permeable AMPARs (CP-AMPARs) through phosphorylation coordinated by AKAP150/PKA/CaN is required along with NMDARs not only for the induction of input-specific LTP and LTD but also for homeostatic plasticity (synaptic scaling) in the hippocampus ^9, 18, 19^.

In spite of the increasing in-depth mechanistic insights into the role of AKAP150 complex in hippocampal-related learning and memory processes, less is known about the normal and pathological roles of AKAP150 complex-dependent signaling in neural processes within reward-related brain circuits that could contribute to reward-related behaviors as well as in the development of neurological and neuropsychiatric illnesses. This is an important area of psychiatric research as human studies of polymorphisms of AKAP5 also indicate that individuals carrying AKAP5 polymorphisms show altered emotional processing and behavioral responses including aggression, expression of anger and impulsivity associated with alterations in the function in limbic regions ^20–22^. Moreover, copy number variations in AKAP5 have been found in DNA samples of schizophrenia patients but not in control subjects ^23^, suggesting the possible involvement of AKAP5 in the pathogenesis of schizophrenia, a neurodevelopmental disorder also linked to reward circuit dysfunction and high rates of addiction ^24–26^. Consistently, in recent years a few studies highlighted the importance of AKAP150 signaling within the ventral tegmental area (VTA)^8, 27–29^, nucleus accumbens (NAc)^30–32^, amygdala^33, 34^ and periaqueductal gray (PAG)^15^ in regulation of synaptic plasticity and the development of depressive states, aversive and drug-related behaviors.

Here, we attempted to address the potential impact of AKAP150-anchored PKA on the lateral habenula (LHb); an anti-reward brain region hub that regulates midbrain monoaminergic centers and is involved in reward/motivation, mood regulation and decision-making. Accumulating evidence indicate that LHb hyperactivity plays an instrumental role in pathophysiology of depression and possibly other mood disorders and substance use disorders, thus LHb is gaining interest as a potential target for neuromodulation and antidepressants^35–38^. LHb neurons are excited by aversive and unpleasant events or the absence of expected reward, and inhibited by unexpected reward, encoding behavioral avoidance and reward prediction errors through suppression of VTA dopamine (DA) and dorsal raphe nucleus (DRN) serotonin systems ^35, 39^. The majority of LHb neurons are believed to be glutamatergic and long-range projecting, although local glutamatergic and GABAergic connections within the LHb are reported ^40–43^. LHb neurons receive glutamatergic, GABAergic and co-releasing glutamate/GABA inputs from the basal ganglia and diverse limbic areas including medial prefrontal cortex (mPFC), entopeduncular nucleus (EP), lateral preoptic area (LPO), lateral hypothalamus (LH), ventral pallidum (VP), medial and lateral septum, central amygdala (CeA), bed nucleus of stria terminals (BNST) as well as receiving reciprocal inputs from the VTA and periaqueductal gray (PAG). LHb projects to the substantia nigra, VTA, rostromedial tegmental area (RMTg), DRN, locus coeruleus (LC) and PAG ^35, 44^. The majority of the glutamatergic output of LHb exerts a potent feedforward inhibitory influence on monoaminergic systems including VTA DA neuronal activity by excitation of GABAergic interneurons and of GABAergic neurons of the RMTg^45–48^.

LHb hyperactivity is found to be a common finding associated with anhedonia, lack of motivation and social withdrawal which reflect some of the core features of reward deficits seen in clinical depression ^36, 49–51^. In general, LHb dysfunction can mediate negative affective states, social deficits, risky decision-making and impulsivity (as shown in patients with depression, schizophrenia, Parkinson’s disease and attention-deficit hyperactivity disorder, ADHD) ^36, 49–57^. Given the known postsynaptic PKA-mediated control of LHb synaptic function and intrinsic excitability by neuromodulatory actions of CRF-CRFR1 signaling^58^, here we investigated the potential impact of AKAP150-anchored PKA on LHb synaptic and neuronal function and CRF neuromodulation within the LHb using whole-cell patch clamp recordings and AKAP150ΔPKA knockin mouse model (hereafter referred to as ΔPKA mice)^14^. The ΔPKA mice have an internal deletion of ten amino acids within the PKA-RII subunit binding domain near the AKAP C terminus. For AKAP150 complex-related studies, they are advantageous compared to AKAP150 knockout mice as the mutation only affects AKAP150-anchoring of PKA without disrupting other AKAP150 interactions^14^. We found that the genetic disruption of PKA anchoring to AKAP150 significantly altered both AMPAR- and GABA_A_R-mediated synaptic transmission and impaired the induction of an eCB-mediated LTD in LHb neurons. Moreover, we observed that ΔPKA mutation enhanced LHb intrinsic excitability which then blunted the excitatory effects of CRF on LHb neuronal activity. Given the significant and multifaceted impact of AKAP150 anchoring of PKA in regulation of glutamatergic transmission and plasticity and neuronal excitability of LHb as well as alteration of CRF regulation of LHb excitability, our data suggest novel roles for AKAP150 complex in normal LHb function and potential contributions of defective AKAP150-mediated PKA anchoring to aberrant LHb activity, dysregulation of CRF neuromodulation within LHb circuits, and hence mood dysregulation.

## 2. Methods

### 2.1. Animals

All experiments were carried out using 5-7wk old male WT (C57/Bl6) and ΔPKA mice in accordance with the National Institutes of Health (NIH) *Guide for the Care and Use of Laboratory Animals* and were approved by the Uniformed Services University and the University of Colorado (Denver) Institutional Animal Care and Use Committees. Mice were group housed in standard cages under a 12hr/12hr light-dark cycle with standard laboratory lighting conditions (lights on, 0600-1800), with ad libitum access to food and water. All procedures were conducted beginning 2–4hr after the start of the light-cycle. All efforts were made to minimize animal suffering and reduce the number of animals used throughout this study.

### 2.2. Slice Preparation

For all electrophysiology experiments several separate cohorts of WT/ ΔPKA mice were used. All mice were anesthetized with isoflurane, decapitated and brains were quickly dissected and placed into ice-cold artificial cerebrospinal fluid (ACSF) containing (in mM): 126 NaCl, 21.4 NaHCO_3_, 2.5 KCl, 1.2 NaH_2_PO_4_, 2.4 CaCl_2_, 1.00 MgSO_4_, 11.1 glucose, 0.4 ascorbic acid; saturated with 95% O_2_-5% CO_2_ as previously described ^59^. Sagittal slices containing LHb were cut at 220µm using a vibratome (Leica; Wetzler, Germany) and incubated in above prepared ACSF at 34°C for at least 1 hour prior to electrophysiological experiments. Slices were then transferred to a recording chamber and perfused with ascorbic-acid free ACSF at 28^°C^.

### 2.3. Electrophysiology

All whole-cell recordings were performed on LHb-containing slices using patch pipettes (3-6 MOhms) and a patch amplifier (MultiClamp 700B) under infrared-differential interference contrast microscopy. Data acquisition and analysis were carried out using DigiData 1440A, pCLAMP 10 (Molecular Devices), Clampfit, Origin 2016 (OriginLab), Mini Analysis 6.0.3 (Synaptosoft Inc.) and GraphPad Prism 10. Signals were filtered at 3 kHz and digitized at 10 kHz. In all of our recordings, the cell input resistance and series resistance were monitored through the experiment and if these values changed by more than 10%, data were not included.

Whole-cell recordings of AMPAR-mediated miniature excitatory postsynaptic currents (mEPSCs) were isolated in ACSF perfused with GABA_A_R antagonist picrotoxin (100µM, Tocris-1128), NMDAR antagonist D-(-)-2-Amino-5-phosphonopentanoic acid (D-APV 50µM, Tocris-0106) and tetrodotoxin (TTX, 1 μM Tocris-1078) and internal solution containing 117 mM Cesium-gluconate, 2.8 mM NaCl, 5 mM MgCl_2_, 2 mM ATP-Na^+^, 0.3 mM GTP-Na^+^, 0.6 mM EGTA, and 20 mM HEPES (pH 7.28, 275–280 mOsm). Whole-cell recordings of GABA_A_R-mediated miniature inhibitory postsynaptic currents (mIPSCs) were isolated in ACSF perfused with the AMPAR antagonist 6,7 dinitroquinoxaline-2,3-dione di-sodium salt (DNQX; 10 μM Tocris-2312/10), strychnine (1 μM Tocris-2785) and tetrodotoxin (TTX, 1 μM). Patch pipettes were filled with 125 mM KCl, 2.8 mM NaCl, 2 mM MgCl_2_, 2 mM ATP-Na^+^, 0.3 mM GTP-Na^+^, 0.6 mM EGTA, and 10 mM HEPES (pH 7.28, 275–280 mOsm). For both mIPSCs and mEPSCs, LHb neurons were voltage-clamped at −70 mV and recorded over 10 sweeps, each lasting 50 seconds.

In some experiments, electrically-evoked AMPAR-mediated EPSCs were isolated and recorded using ACSF containing picrotoxin (100 μM). The patch pipettes were filled with cesium-gluconate based solution as described above for mEPSC recordings. Cells were voltage-clamped at −70 mV, except during LTD protocol. Paired AMPAR-mediated EPSCs were stimulated at 0.1 Hz (100 ms) using a bipolar stainless steel stimulating electrode placed ∼200-400 µm from the recording site in stria medularis in LHb slices. The stimulation intensity was adjusted so that the amplitude of synaptic responses ranged about ∼50% of maximum response. LTD was induced using low frequency stimulation, LFS, 1 Hz for 15 min while LHb neurons were voltage-clamped at −40 mV.

To assess LHb intrinsic excitability and membrane properties, LHb slices were perfused with ascorbic-free ACSF and patched with potassium gluconate-based internal solution (130 mM K-gluconate, 15 mM KCl, 4 mM ATP-Na^+^, 0.3 mM GTP-Na^+^, 1 mM EGTA, and 5 mM HEPES, pH adjusted to 7.28 with KOH, osmolarity adjusted to 275 to 280 mOsm). LHb intrinsic excitability experiments were performed with fast-synaptic transmission blockade by adding DNQX (10µM), picrotoxin (100 µM), and D-APV (50 µM) to the ACSF. LHb neurons were given increasingly depolarizing current steps at +10pA intervals ranging from +10pA to +100pA, allowing us to measure AP generation in response to membrane depolarization (5 sec duration). Current injections were separated by a 20s interstimulus interval and neurons were kept at ∼-65 to −70 mV with manual direct current injection between pulses. Resting membrane potential (RMP) was assessed immediately after achieving whole-cell patch configuration in current clamp mode. Input resistance (Rin) was measured during a −50pA step (5s duration) and calculated by dividing the steady-state voltage response by the current-pulse amplitude (−50pA) and presented as MOhms (MΩ). The number of APs induced by depolarization at each intensity was counted and averaged for each experimental group. As previously described^58^ AP number, AP threshold, fast and medium after-hyperpolarization amplitudes (fAHP and mAHP), AP halfwidth, AP amplitude were assessed using Clampfit and measured at the current step that was sufficient to generate the first AP/s.

### 2.4. Drugs

For all drug experiments a within-subjects experimental design was employed. Stock solutions for CRF were prepared in distilled water and diluted (1:1000) to final concentration in ACSF of 250nM. Baseline recordings were first performed (depolarization-induced AP/mIPSC/mEPSC) for each neuron and then CRF (250 nM Tocris-1151) was added to the slice by the perfusate and response tested following 30-45min of CRF application.

### 2.5. Immunohistochemistry

Mice were anesthetized with an intraperitoneal injection containing ketamine (85 mg/kg) and xylazine (10 mg/kg) and perfused through the aorta with 200 ml of 1x phosphate buffered saline (PBS), followed by 250 ml of 4% paraformaldehyde (PFA) (Santa Cruz). The brains were dissected and placed in 4% PFA for 24 h and then cryoprotected by submersion in 20% sucrose for 3 d, frozen on dry ice, and stored at-70°C until sectioned. Sections of the LHb were cut using a cryostat (Leica CM1900) and mounted on slides. Serial coronal sections (20 μm) of the midbrain containing the LHb (from −2.64 to −4.36 mm caudal to bregma (Paxinos and Watson, 2007) were fixed in 4% PFA for 5 min, washed in 1x PBS, and then blocked in 10% normal goal serum (NGS) containing 0.3% Triton X-100 in 1x PBS for 1 h. Sections were incubated in goat anti-AKAP150 antibody (1:500, Santa Cruz Sc-6445) in carrier solution (5 % NGS in 0.1% Triton X-100 in 1x PBS) overnight at room temperature. After rinsing in 1x PBS, sections were incubated for 2 h in Alexa Fluor® 488 labeled chicken anti-goat IgG (diluted 1:200). Finally, sections were rinsed in 1x PBS, dried, and cover slipped with Prolong mounting medium containing DAPI to permit visualization of nuclei. Background staining was assessed by omission of primary antibody in the immunolabeling procedure (negative control). Brain tissue sections of mouse with previously established presence of AKAP150 immunoreactive neurons (hippocampus, VTA) were also processed as positive control tissues. Images were captured using a Leica DMRXA Fluorescence microscope.

### 2.6. Data analysis

Values are presented as means ± SEM. Statistical significance was determined using unpaired or paired two tailed Student’s t-test, two-way ANOVA or repeated-measures ANOVA (RM-ANOVA) with Bonferroni post hoc analysis. The threshold for significance was set at *P < 0.05 for all analyses. The peak values of the evoked paired EPSCs were measured relative to the same baseline. A stable baseline value was considered in each sweep of paired pulses starting at 20-50ms right before the emergence of the EPSC current using p-Clamp 10 software. The paired pulse ratio (PPR) was calculated as the amplitude of the second EPSP divided by the amplitude of the first EPSC. The inverse square of the coefficient of variation (CV=SD/mean) was also used as the second measure for identifying the presynaptic expression of plasticity. For calculating significance of EPSC amplitude changes after LTD induction protocol, amplitudes of EPSCs to the first pulse were used. Mini Analysis software was used to detect and measure mIPSCs and mEPSCs using preset detection parameters of mIPSCs and mEPSCs with an amplitude cutoff of 5 pA. The Kolmogorov–Smirnov test (KS test) was performed for the statistical analyses of cumulative probability plots of mEPSCs and mIPSCs. All statistical analyses were performed using GraphPad Prism 10.

## 3. Results

### 3.1. AKAP150 expression in the LHb

Figure 1 depicts a representative 40x image of LHb of a young adult male mouse taken at AP location (−1.34 relative to bregma). We observed wide expression of AKAP150 in the LHb at three AP locations (−1.06, −1.34 and −1.46) in 4 wild type (WT) mice.

**Figure 1:**
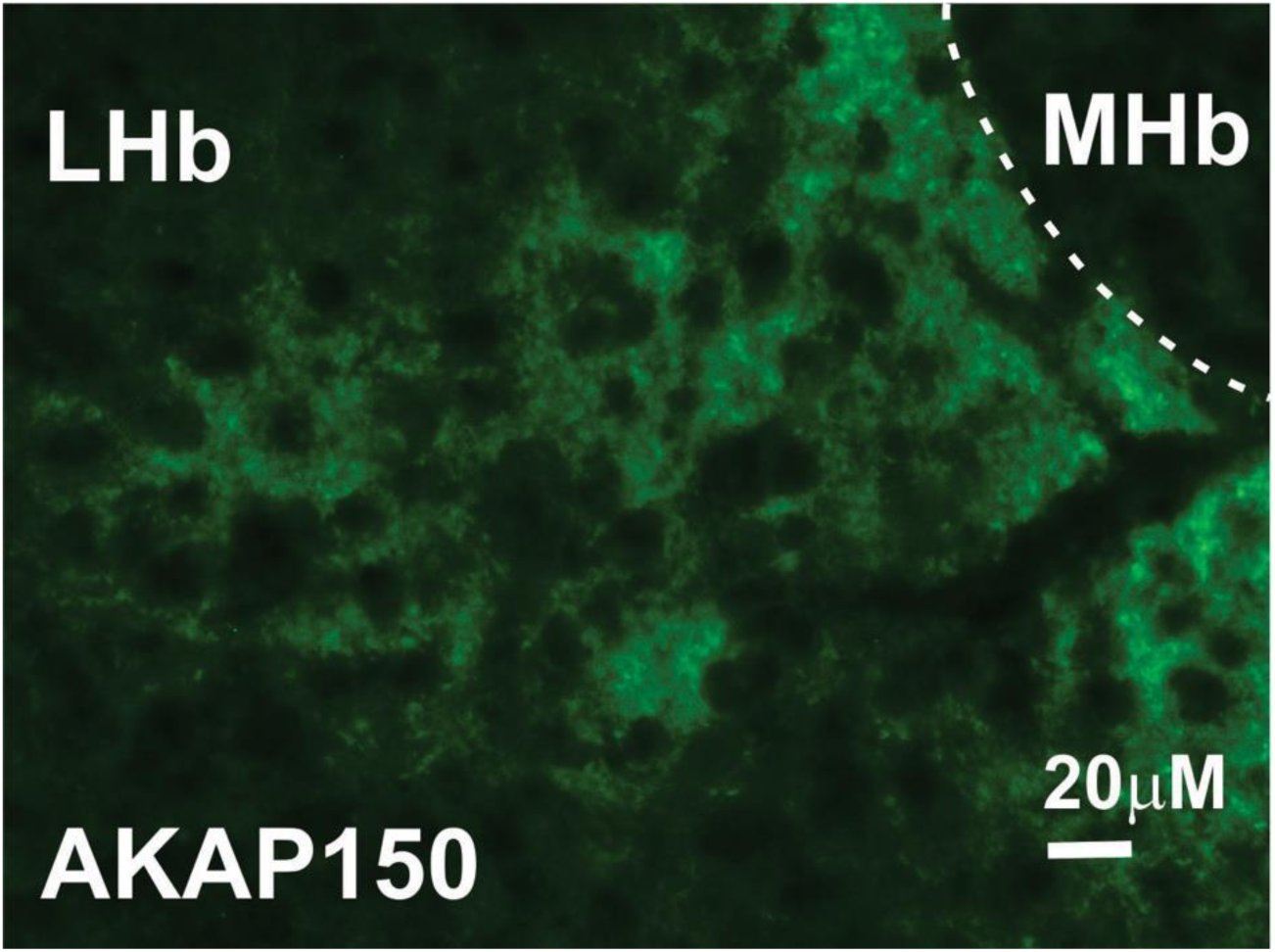
AKAP 150 is expressed in the LHb. Example of a brain section stained with antibody to AKAP150 (green), showing the expression of AKAP150 in the mouse LHb. Scale bar, 20μm.

### 3.2. Effects of AKAP-ΔPKA mutation on synaptic transmission and glutamatergic LTD in LHb neurons

To examine the effects of genetic disruption of PKA anchoring to AKAP150 on AMPAR and GABA_A_R-mediated synaptic transmission, we recorded either mEPSCs (Figure 2) or mIPSCs (Figure 3) from LHb neurons from WT and and ΔPKA mice with deficiency in PKA-anchoring to AKAP150 ^14^. ΔPKA mutation significantly decreased the average amplitude (inset in Figure 2B), frequency (inset in Figure 2C) and charge transfer (inset in Figure 2D) of mEPSCs and correspondingly shifted the cumulative probability curves of mEPSC amplitude (to the left indicative of smaller amplitude, Figure 2B), inter-event interval (IEI, to the right indicative of lower frequency, Figure 2C) and charge transfer (to the left indicative of lower charge transfer, Figure 2D) without altering mEPSC tau decay (Figure 2, unpaired Student’s t-tests, Kolmogorov-Smirnov tests, p<0.05, p<0.01, p<0.001, p<0.0001), suggesting both pre- and postsynaptic suppression of glutamatergic transmission in LHb neurons. Only the cumulative probability curve of mIPSC amplitude (Figure 3B) was significantly shifted to the right by this genetic AKAP-PKA disruption, which may indicate an increase in postsynaptic GABA_A_R function at a subset of GABAergic synapses onto LHb neurons (Figure 3, Kolmogorov-Smirnov tests, p<0.0001). All other mIPSC properties were not significantly different between ΔPKA and WT.

**Figure 2.**
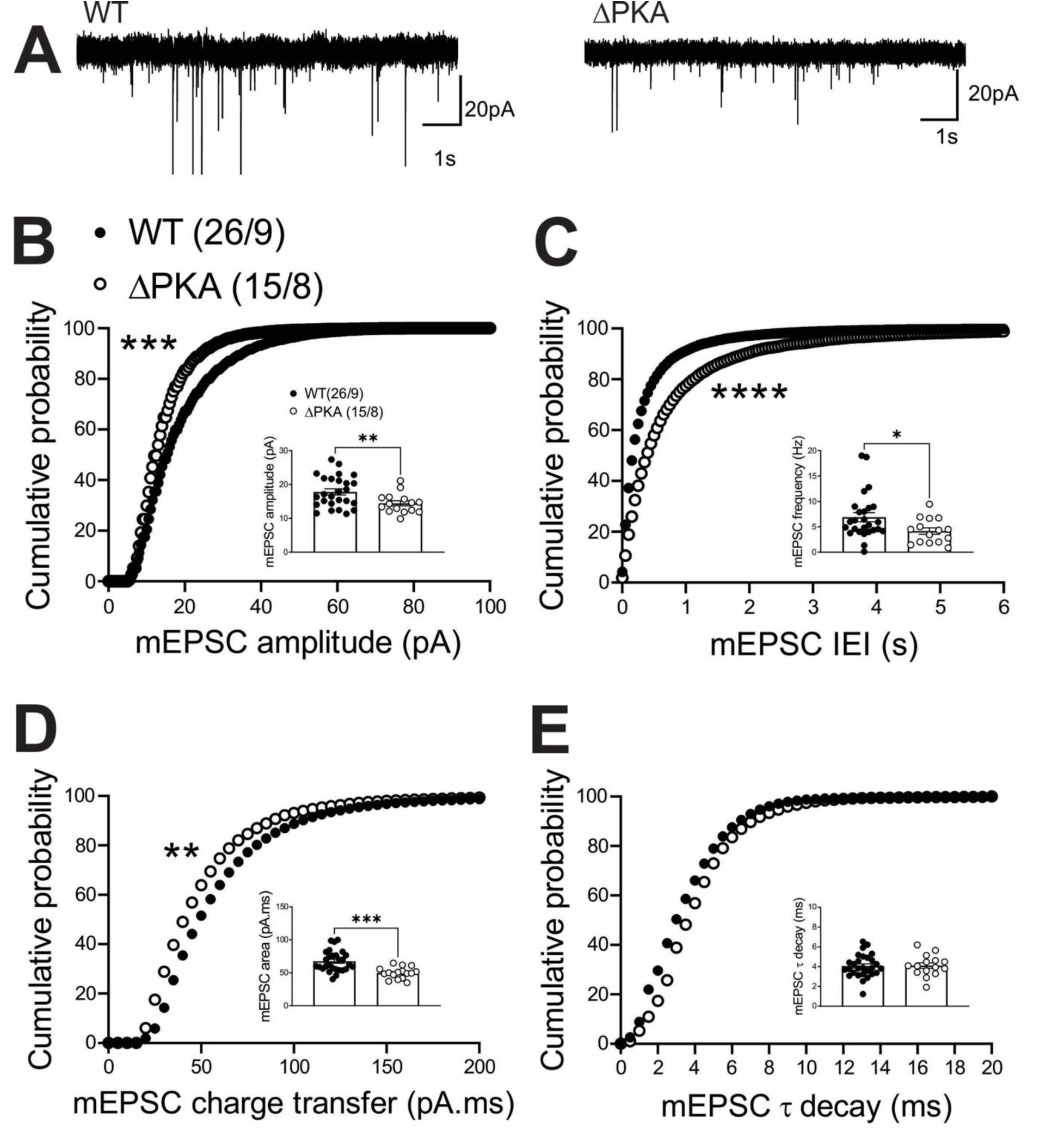
Genetic disruption of AKAP150-anchored PKA depressed glutamatergic transmission in LHb neurons. **(A)** Sample AMPAR-mediated mEPSC traces from WT (left) and ΔPKA mice (calibration bars: 20 pA/1s). Average bar graphs of mEPSC and cumulative probability plots of **(B)** amplitude, **(C)** frequency (inter-event interval), **(D)** charge transfer and **(E)** τ decay for all mEPSCs in WT (filled symbols) and ΔPKA (open symbols) mice (n=15-26 cells from 8-9 mice/group).

**Figure 3.**
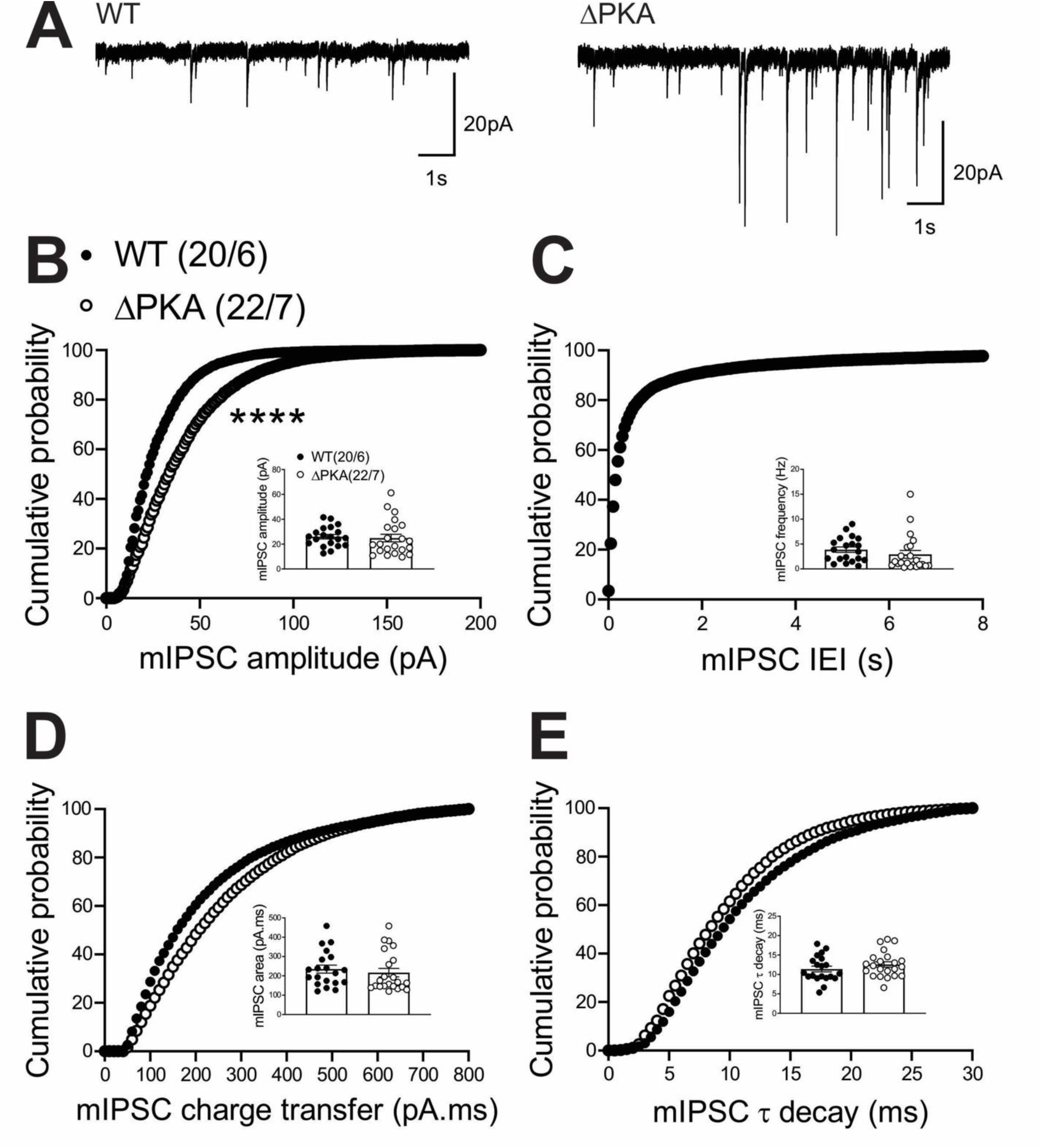
Genetic disruption of AKAP150-anchored PKA potentiated GABAergic transmission in LHb neurons. **(A)** Sample GABA_A_R-mediated mIPSC traces from WT (left) and ΔPKA mice (calibration bars: 20 pA/1s). Average bar graphs of mIPSC and cumulative probability plots of **(B)** amplitude, **(C)** frequency (inter-event interval), **(D)** charge transfer and **(E)** τ decay for all mIPSCs in WT (filled symbols) and ΔPKA (open symbols) mice (n=15-26 cells from 8-9 mice/group).

Previously, it has been reported that LFS can induce a retrograde presynaptic eCB-mediated LTD of the AMPAR-mediated electrically-evoked EPSCs in LHb neurons by postsynaptic activation of group I metabotropic glutamate receptors (mGluR-LTD)^60^ or through calcium permeable AMPARs (CP-AMPARs) that further activate NMDARs ^61,62^. Here, we also used an identical LFS protocol to induce a presynaptic LTD and assessed PPRs and 1/CV^2^ values as the two main indicators of presynaptic expression of synaptic plasticity. As shown in Figure 4, the LTD protocol strongly induced eCB-LTD onto LHb neurons in WT mice (Figure 4A and 4C) which was associated with significant increases in PPRs (Figure 4D) and corresponding decreases in 1/CV^2^ (Figure 4E) suggesting the expression of eCB-mediated LTD. On the other hand, LHb neurons from ΔPKA mice (Figure 4B and 4C) were unable to express this presynaptic glutamatergic LTD (Figure 4, LTD: WT, F (1.45, 6.30) = 17.27; ΔPKA: F (1.22, 3.67) = 1.083, p=0.38, RM ANOVA. PPRs and I/CV^2^: unpaired Student’s t-test, p <0.01, p<0.05).

**Figure 4:**
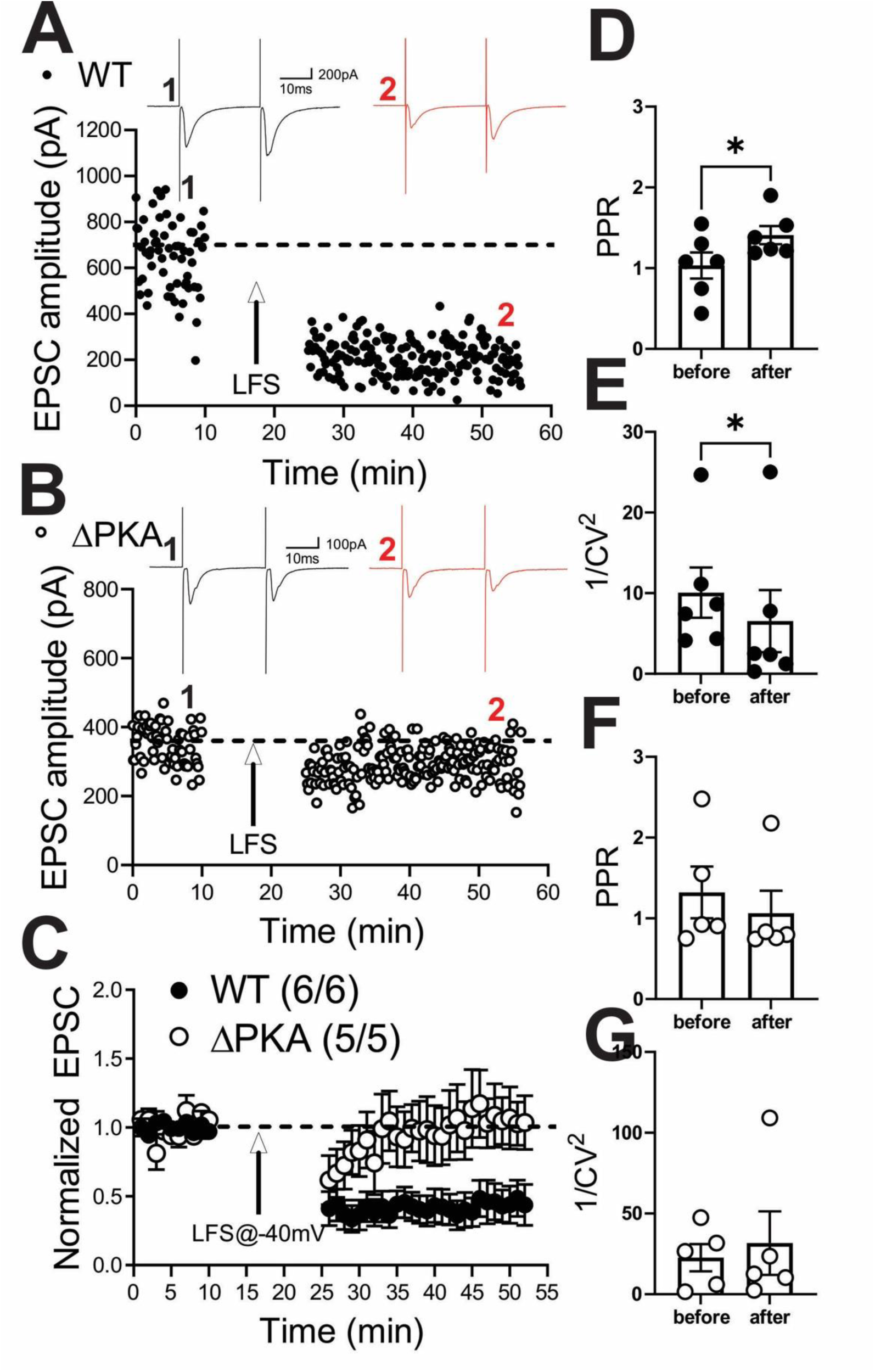
Genetic disruption of AKAP150-anchored PKA impaired eCB-LTD induction in LHb neurons. **(A)** and **(B)** Single experiments showing induction of LTD recorded in LHb neurons from WT **(A)** and ΔPKA **(B)** mice. At the arrow, LTD was induced using LFS while cells were depolarized at −40mV. Insets: averaged EPSCs before (black, labeled as **1**) and 25 min after LFS (red, labeled as **2**). In this and all figures, ten consecutive traces from each condition were averaged for illustration as inset. Calibration: 100-200 pA, 10 ms. **(C)** Average LTD experiments with corresponding PPRs **(D** and **F)** and 1/CV^2^ **(E** and **G)** in LHb neurons recorded from WT (filled symbols) and ΔPKA (open symbols) neurons (n=5-6 cells/5-6 mice/group).

### 3.3. Effects of AKAP-ΔPKA mutation on LHb intrinsic excitability

Given that PKA regulation of various ion channels in the neuronal membrane requires anchoring of PKA by AKAP150^63^, it was possible that the genetic disruption of PKA-AKAP association in ΔPKA mice could also impact intrinsic plasticity through changes in trafficking and/or function of a number of voltage-gated channels. Consistently, we observed that LHb neurons of ΔPKA mice exhibited significantly higher intrinsic excitability in the absence of synaptic transmission compared to those from WT mice (Figure 5A). Furthermore, ΔPKA mutation-induced intrinsic plasticity was associated with lower amplitude of mAHPs (Figure 5D), and shorter AP half widths (Figure 5G) suggesting that the genetic disruption of AKAP150-PKA anchoring modified intrinsic active and passive neuronal membrane properties, which could also influence synaptic conductance (Figure 5 A-G, intrinsic excitability: F (1, 209) = 21.22, 2-way ANOVA; mAHPs and AP half width: unpaired Student’s t test, p<0.05, p<0.01, p<0.0001).

**Figure 5:**
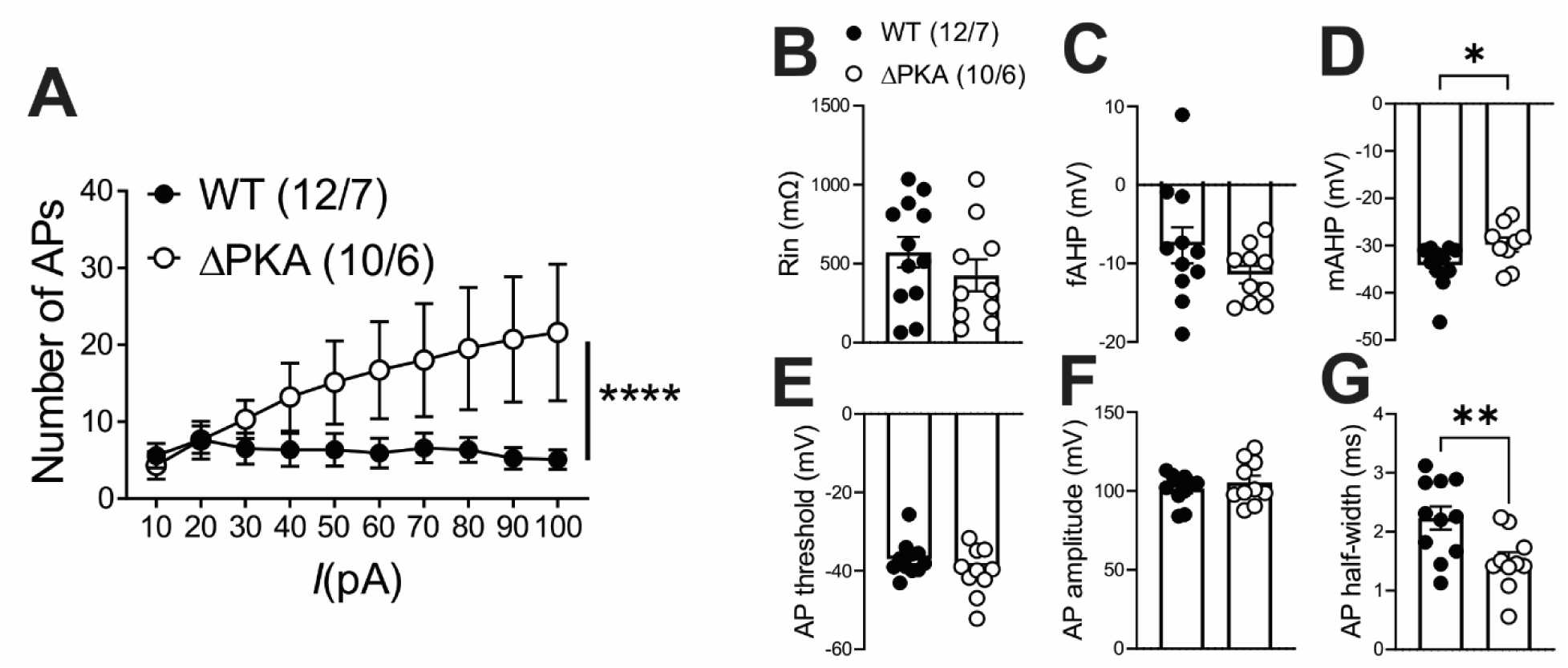
Genetic disruption of AKAP150-anchored PKA increased LHb intrinsic excitability and occluded the effects of CRF on LHb excitability. All recordings in this graph were performed with fast synaptic transmission blocked. **(A-G)** AP recordings in response to depolarizing current steps and corresponding measurements of Rin, fAHP, mAHP, AP threshold, AP amplitude and AP half width in LHb neurons from WT (black filled symbols) and ΔPKA (black open symbols) (n=10-12 cells/ 6-7 mice/group).

### 3.4. Effects of AKAP-ΔPKA mutation on CRF neuromodulation within the LHb

Previously, we demonstrated that the LHb is a highly CRF-responsive brain region with PKA-dependent regulation of LHb synaptic inhibition and intrinsic excitability. We showed that CRF acting through postsynaptic CRFR1 and PKA signaling increases LHb excitability through PKA-dependent suppression of small conductance potassium SK channel activity, as well as presynaptic GABA release via retrograde eCB-CB1 receptor (CB1R) signaling in rat LHb neurons without any significant alterations in glutamatergic transmission^58^. In contrast to our earlier findings in rat LHb where exogenous CRF did not alter mEPSCs, CRF bath application significantly decreased the average frequency of mEPSCs (inset in Figure 6C) and correspondingly shifted the cumulative probability curves of mEPSC IEI to the right (Figure 6C) (Figure 6, paired Student’s t-tests, Kolmogorov-Smirnov tests, p<0.05, p<0.0001), suggesting of CRF-induced suppression of presynaptic glutamate release in mouse LHb neurons. This diminishing effect of CRF on presynaptic glutamate release was absent in LHb neurons of ΔPKA mice and we even detected a slight but significant leftward shift in the cumulative probability curves of mEPSC IEI (Figure 7C) that may suggest an unmasked potentiating effects of CRF on presynaptic glutamate release upon defective AKAP150-PKA associations (Figure 7, Kolmogorov-Smirnov tests, p<0.05).

**Figure 6.**
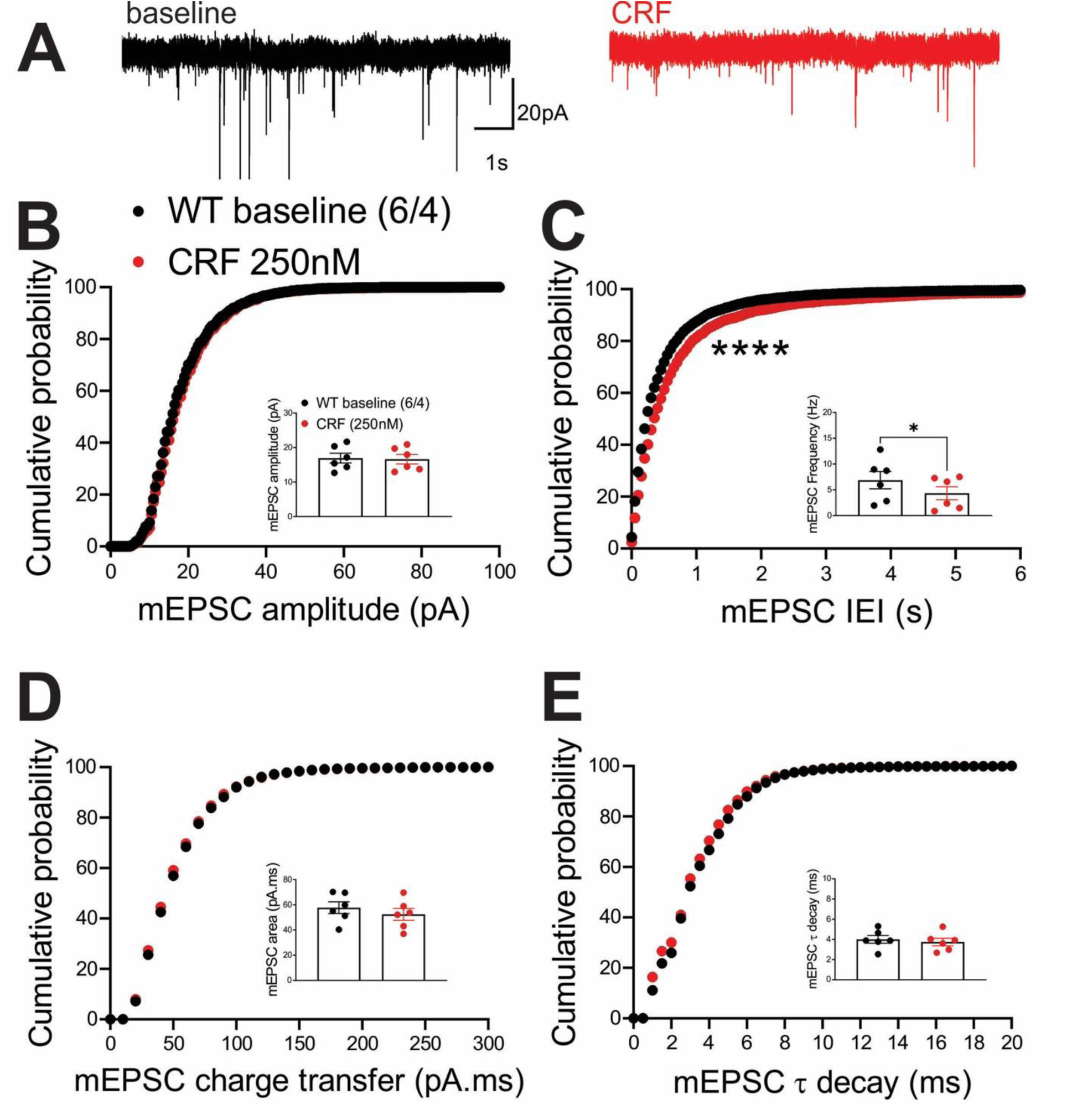
CRF decreased presynaptic glutamate release in the LHb of WT mice. **(A)** Sample AMPAR-mediated mEPSC traces from WT mouse before (black) and after CRF application (red, 250nM) (calibration bars: 20 pA/1s). Average bar graphs of mEPSC and cumulative probability plots of **(B)** amplitude, **(C)** frequency (inter-event interval), **(D)** charge transfer and **(E)** τ decay for all mEPSCs in WT mice before (black filled symbols) and after CRF (red filled symbols) (n=6 cells from 4 mice).

**Figure 7.**
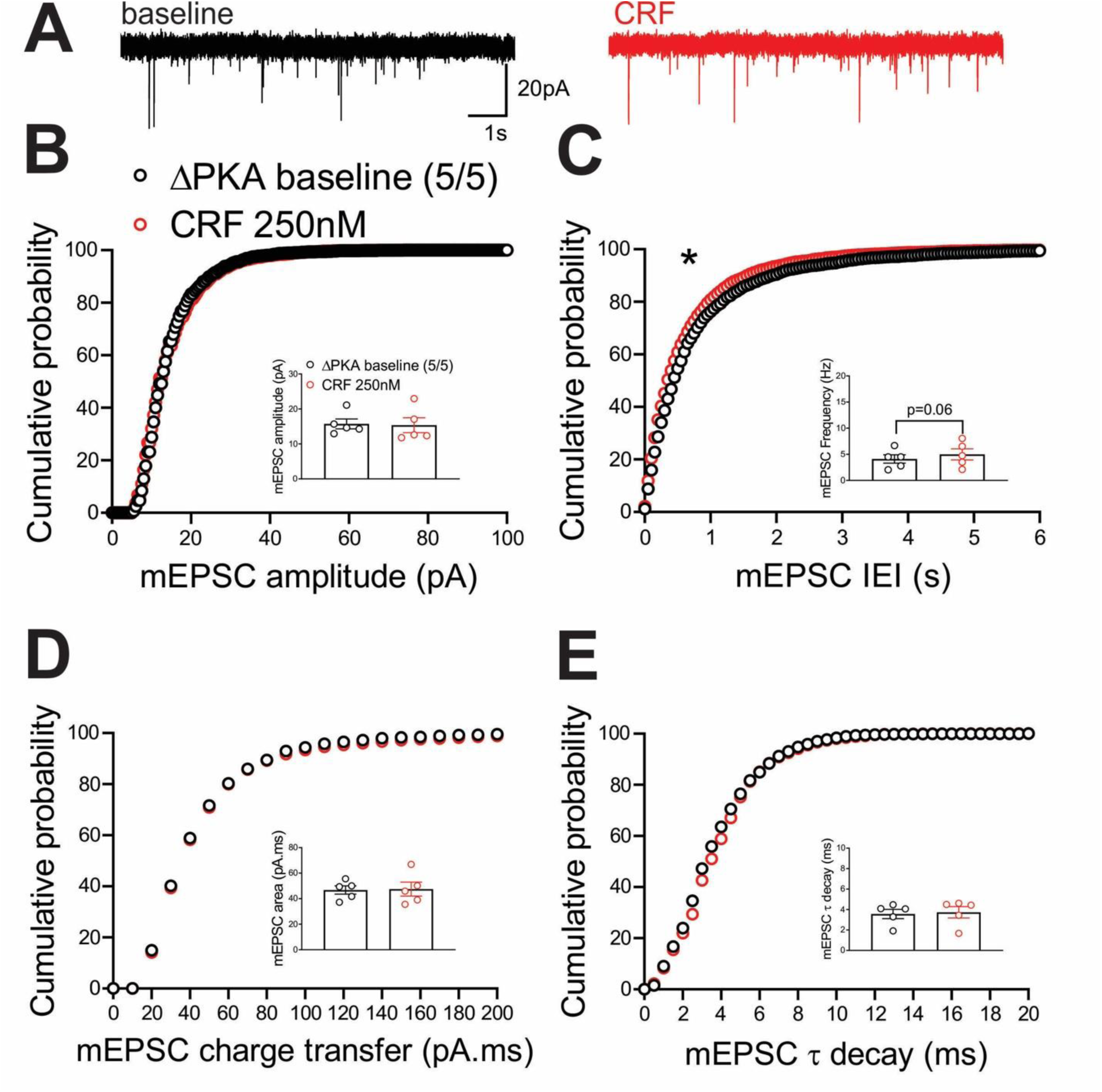
CRF slightly potentiated presynaptic glutamate release in the LHb of ΔPKA mice. **(A)** Sample AMPAR-mediated mEPSC traces from ΔPKA mouse before (black) and after CRF application (red, 250nM) (calibration bars: 20 pA/1s). Average bar graphs of mEPSC and cumulative probability plots of **(B)** amplitude, **(C)** frequency (inter-event interval), **(D)** charge transfer and **(E)** τ decay for all mEPSCs in ΔPKA mice before (black open symbols) and after CRF (red open symbols) (n=5 cells from 5 mice).

In contrast, we observed similar effects of the ΔPKA mutation on GABAergic transmission in mouse LHb and rat LHb; exogenous CRF significantly diminished the average frequency of mIPSCs (inset in Figure 8C) and resulted in a significant shift in the cumulative probability curves of mIPSC IEI to the right (Figure 8C) (Figure 8, paired Student’s t-tests, Kolmogorov-Smirnov tests, p<0.05, p<0.0001), indicating CRF-induced suppression of presynaptic GABA release onto mouse LHb neurons. This diminishing effect of CRF on presynaptic GABAergic transmission remained intact in LHb neurons of ΔPKA mice as evident with a smaller but still significant rightward shift in the cumulative probability curves of mIPSC IEI (Figure 9C). However, CRF additionally decreased the average amplitude (inset in Figure 9B) and charger transfer (inset in Figure 9D) of mIPSCs as well as shifted their corresponding cumulative probability curves of mIPSC amplitude (Figure 9B) and charge transfer (Figure 9D) to the left. These results may indicate that CRF alters postsynaptic function and/or trafficking of GABA_A_Rs upon disruption of AKAP150-PKA association, potentially favoring AKAP150-dependent regulation of anchored PKC and/or CaN phosphatase activity, both of which can negatively regulate postsynaptic GABA_A_Rs ^29, 64^, downstream of CRF-CRFR-PKC signaling^65, 66^ (Figure 9, paired Student’s t-tests, Kolmogorov-Smirnov tests, p<0.05, p<0.0001).

**Figure 8.**
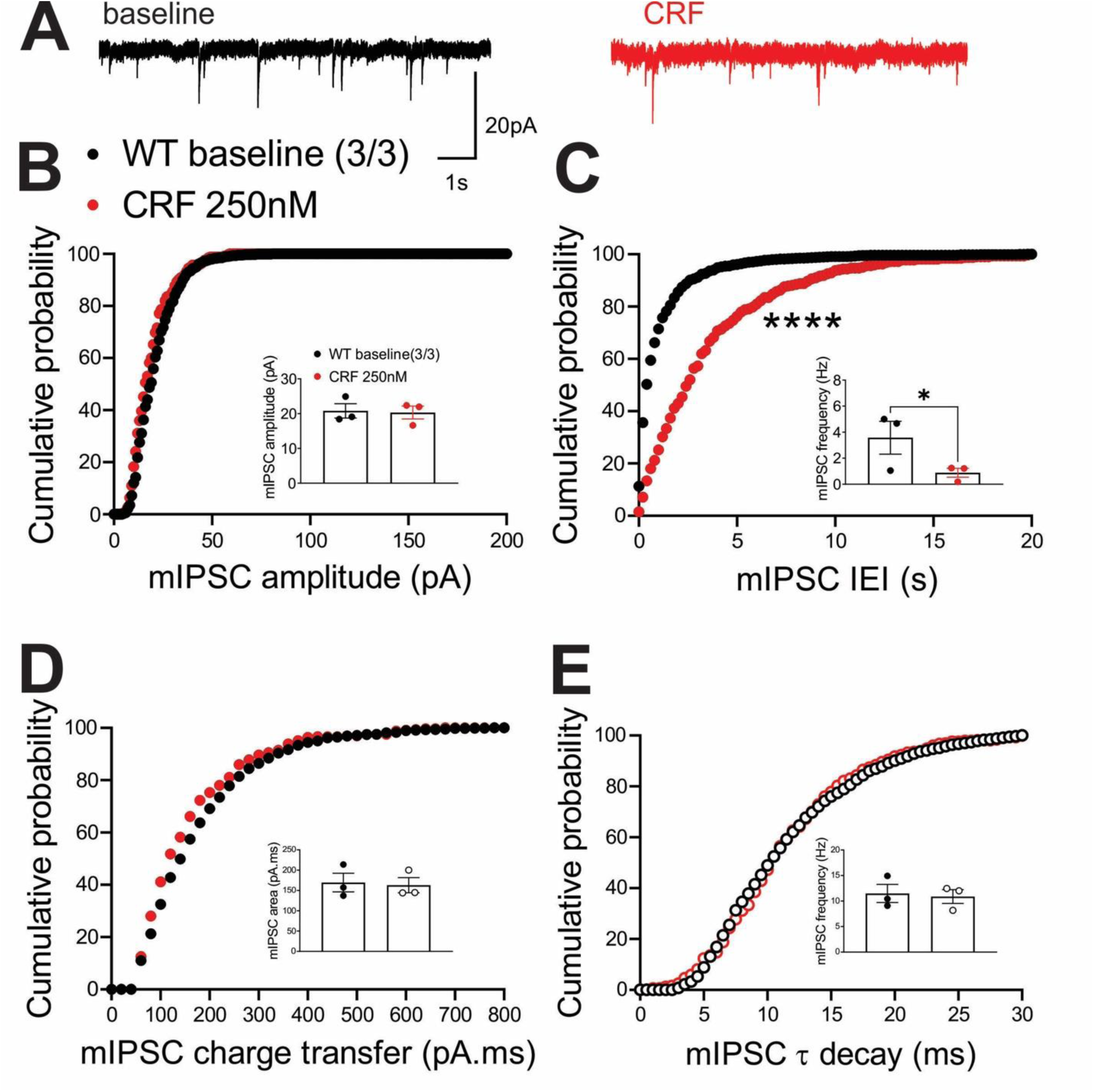
CRF significantly suppressed presynaptic GABA release in the LHb of WT mice. **(A)** Sample GABA_A_R-mediated mIPSC traces from WT mouse before (black) and after CRF application (red, 250nM) (calibration bars: 20 pA/1s). Average bar graphs of mIPSC and cumulative probability plots of **(B)** amplitude, **(C)** frequency (inter-event interval), **(D)** charge transfer and **(E)** τ decay for all mIPSCs in WT mice before (black filled symbols) and after CRF (red filled symbols) (n=3 cells from 3 mice).

**Figure 9.**
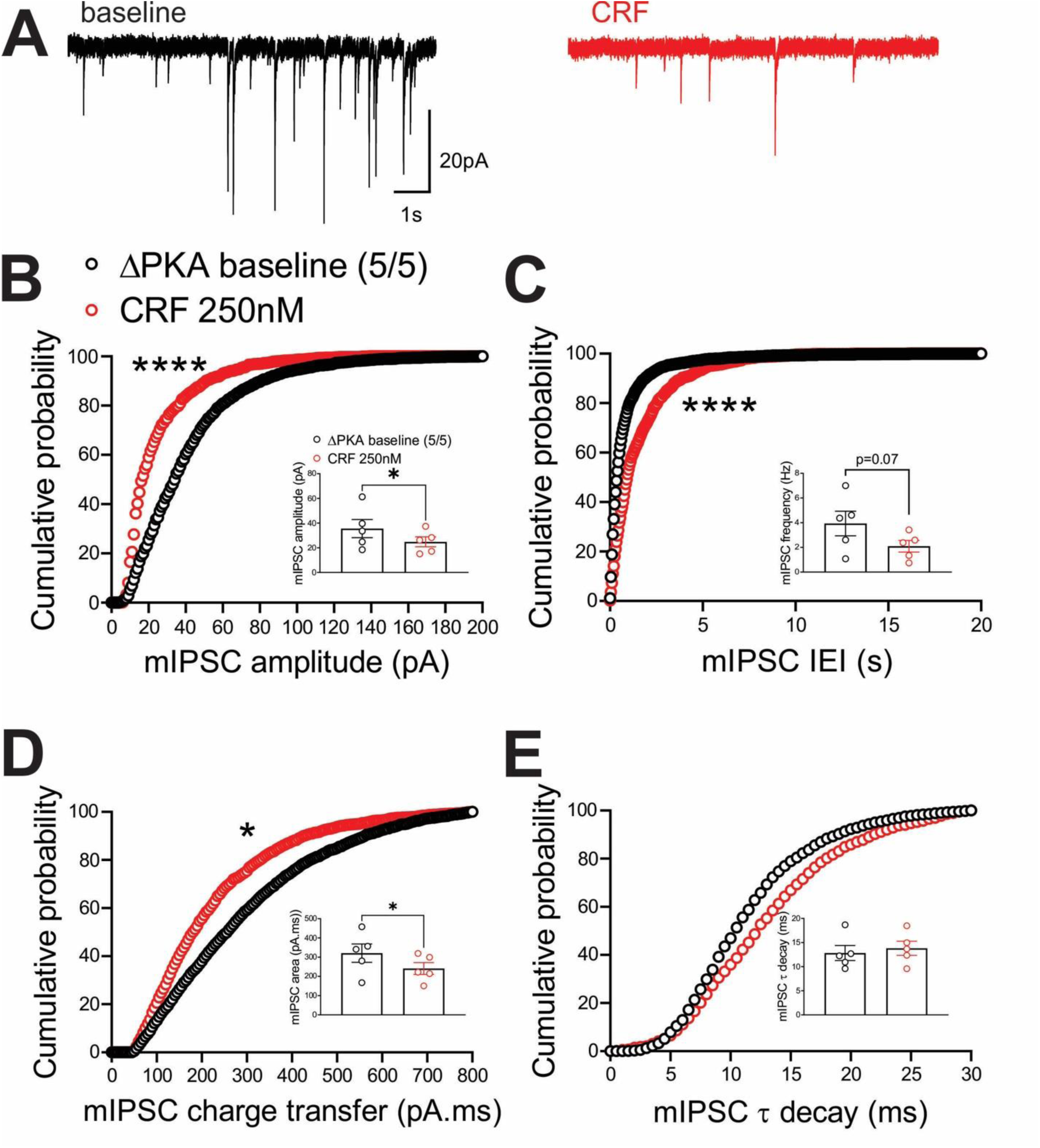
CRF suppressed presynaptic GABA release (to a lesser extent than that of WT) but also postsynaptically depressed GABAergic transmission in the LHb of ΔPKA mice. **(A)** Sample GABA_A_R-mediated mIPSC traces from ΔPKA mouse before (black) and after CRF application (red, 250nM) (calibration bars: 20 pA/1s). Average bar graphs of mIPSC and cumulative probability plots of **(B)** amplitude, **(C)** frequency (inter-event interval), **(D)** charge transfer and **(E)** τ decay for all mIPSCs in ΔPKA mice before (black open symbols) and after CRF (red open symbols) (n=5 cells from 5 mice).

Similar to our earlier findings in rat LHb^58^, CRF bath application exerted similar effects on intrinsic excitability and intrinsic membrane properties of male mouse LHb. We found that CRF significantly increased LHb intrinsic excitability (with blocked fast AMPAR, NMDAR and GABA_A_R-mediated transmission) (Figure 10A-B) coincident with higher input resistance (Figure 10C), reduced levels of mAHPs (Figure 10E), lower AP threshold (Figure 10F) and smaller AP amplitudes (Figure 10G) in LHb neurons. The only exception was that CRF-induced increases in fAHPs were not observed in mouse LHb (Figure10D) (Figure 10 A-H, 2-way RM ANOVA, F (1, 7) = 8.23, p<0.05). On the other hand, we observed that the excitatory actions of CRF in LHb of ΔPKA mice was absent (Figure 10I-J), although CRF was able to increase the amplitude of fAHPs (Figure 10L) in LHb neurons of ΔPKA mice (Figure 10 I-P, 2-way RM ANOVA, F (1, 6) = 0.4436, p=0.53).

**Figure 10:**
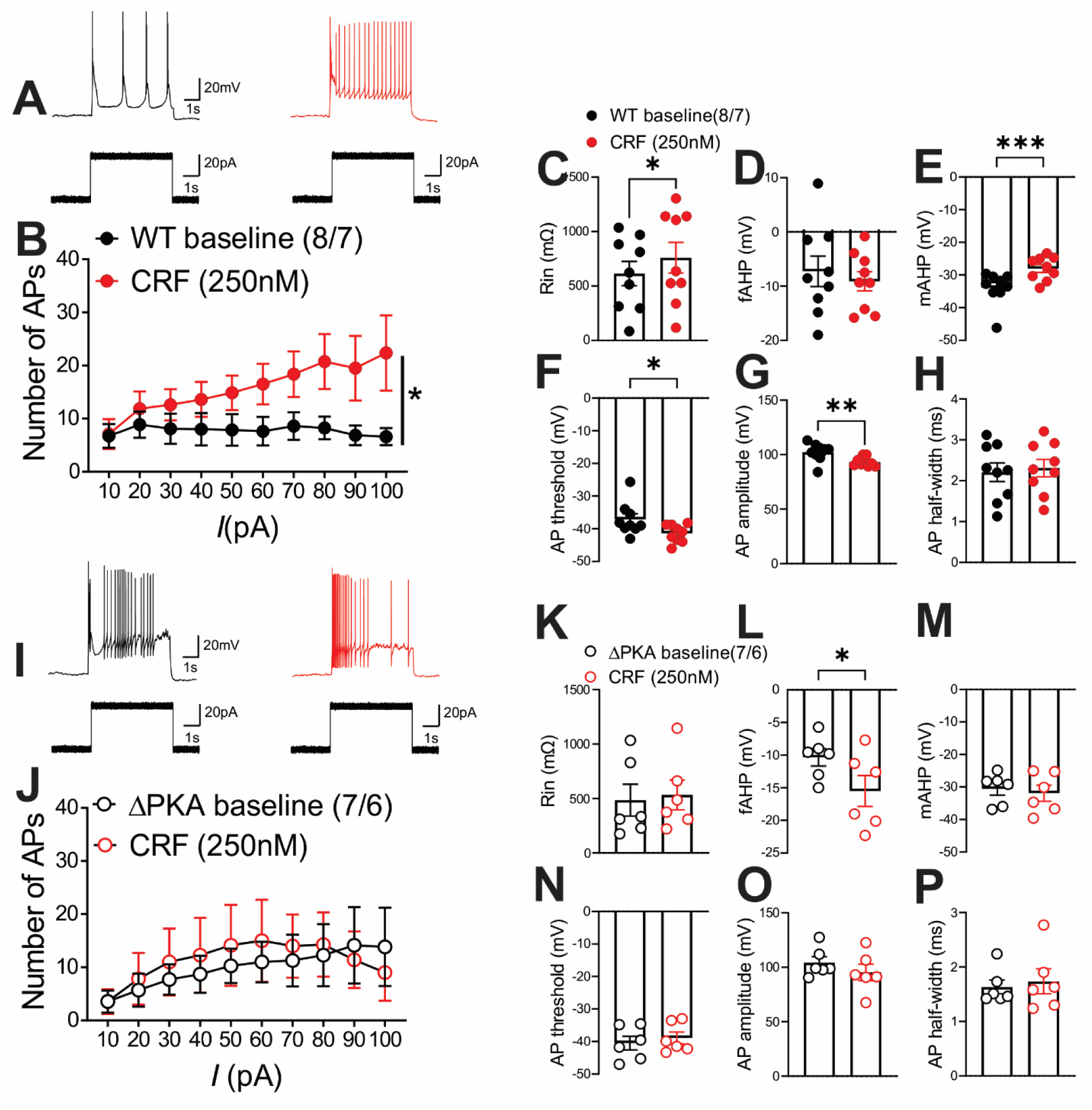
Genetic disruption of AKAP150-anchored PKA occluded the effects of CRF on LHb excitability. All recordings in this graph were performed with fast synaptic transmission blocked. **(A-H)** AP recordings in response to depolarizing current steps with representative AP traces (in response to a 40pA current step) and corresponding measurements of Rin, fAHP, mAHP, AP threshold, AP amplitude and AP half width before (baseline, black filled symbols), after CRF (250nM, red filled symbols) bath application in LHb neurons from WT mice (n=8 cells/7 mice). **(I-P)** AP recordings in response to depolarizing current steps with representative AP traces (in response to a 40pA current step) and corresponding measurements of Rin, fAHP, mAHP, AP threshold, AP amplitude and AP half width before (baseline, black open symbols), after CRF (250nM, red open symbols) bath application in LHb neurons from ΔPKA mice (n=7 cells/6 mice).

## 4. Discussion

Most of our understanding of the role of AKAP150 in the brain relates to hippocampal studies but emerging evidence also suggests novel roles for AKAP150 in reward-related brain regions critical for control of mood, motivation, reward, and stress responses^8, 15,27–31, 33, 34^. Here, we uncovered a multifaceted essential regulatory role of AKAP150 in synaptic function, neuronal activity and CRF neuromodulatory actions within the LHb using the ΔPKA knockin mouse model with deficiency of AKAP150 anchoring of PKA. We found that in LHb neurons AKAP150-anchored PKA is required for postsynaptic regulation of AMPAR trafficking and/or function at glutamatergic synapses and for the expression of glutamatergic eCB-LTD. In contrast, AKAP150-PKA signaling may provide an inhibitory feedback mechanism for postsynaptic trafficking and/or function of GABA_A_Rs or regulate gene expression programs that indirectly control inhibitory synaptic strength ^67^. Moreover, we found that defects in AKAP150-PKA mediated expression, trafficking, and/or gating of potassium channels that regulate LHb excitability could blunt CRF neuromodulatory effects within the LHb.

Postsynaptic AMPAR and GABA_A_R trafficking and function is necessary for maintaining basal synaptic transmission as well as induction and expression of synaptic plasticity which can be altered through phosphorylation-dephosphorylation processes that require AKAP150^63^. There are four subunits of AMPARs (GluR1–GluR4) of which LHb neurons express high levels of GluA1-containing rectifying AMPARs that lack GluA2 (also called calcium permeable, CP-AMPARs with fast kinetics, high conductance and strong inward rectification) but also express low levels of both GluA2- containing AMPARs (with slower kinetics, low conductance and impermeability to calcium) and NMDARs at their glutamatergic synapses^68, 69^. AKAP150-anchored PKA is shown to phosphorylate Ser-845 on the GluA1 subunit of AMPARs to increase membrane trafficking of AMPARs at glutamatergic synapses, while AKAP150-anchored protein kinase C (PKC), through phosphorylation of Ser-831 on GluA1 results in emergence of CP AMPARs at synapses^70^. In the LHb, it has been shown that cocaine can induce synaptic potentiation and hyperexcitability in LHb neurons projecting to RMTg (LHb^→RMTg^ neurons) through Ser-845 phosphorylation of GluA1 that increases trafficking of GluA1 AMPARs in LHb^→RMTg^ neurons^71^. Additionally, phosphorylation of Ser-831 on GluA1 subunit of AMPARs by β-calcium/calmodulin-dependent kinase type II (CaMKIIβ) in the LHb is shown to promote GluA1 AMPAR insertion into synapses and glutamatergic potentiation, resulting in LHb hyperactivity and behavioral depression^72^. Given that the knockin mutation in ΔPKA mice leads to reductions in postsynaptic PKA localization in dendritic spines ^14^, our observation of lower levels of mEPSC amplitude and charge transfer in ΔPKA mice are most likely due to decreased PKA-dependent Ser-845 phosphorylation of GluA1 rectifying AMPARs by the genetic disruption of AKAP150 anchoring of PKA to GluA1. Note that the decrease in frequency of mEPSC in ΔPKA mice is likely postsynaptic, related to an increase in the number of silent synapses following the loss of AMPARs at the synapse^73^ rather than a change in presynaptic glutamate release. Whether, upregulation of CaMKII-regulated AKAP79/150 depalmitoylation that is important in AKAP150 removal from dendritic spine and structural LTD^74^ is favored in ΔPKA mice is an open question.

Interestingly, disruption of PKA anchoring in ΔPKA mice is shown to impair an NMDAR-dependent LTD induced by prolonged low-frequency stimulation (LFS; 1 Hz, 15 min similar to the LTD protocol in LHb) in CA1 hippocampal neurons of 2-week old mice due to decreased S845 phosphorylation of CP-AMPARs in ΔPKA mice that prevents the AKAP150-PKA-dependent transient recruitment of CP-AMPARs to the synapse that is required for hippocampal LTD induction^9^. Interestingly, LFS can also induce eCB-mediated LTD in the LHb through increased activity of CP-AMPAR (as a major source of calcium) that further engages NMDARs to trigger eCB-LTD ^61, 62^. Given that the majority of AMPARs in the LHb are CP-AMPARs, a reduced level of CP-AMPARs in LHb neurons of ΔPKA mice could result in lower levels of depolarization and postsynaptic calcium needed for eCB production, thereby deficits in induction and expression of eCB-LTD. Moreover, since basal PKA phosphorylation of L-type calcium channels (LTCC) is necessary for depolarization-induced activation of LTCCs, the ΔPKA mutation could further diminish Ca^2+^ influx through LTCC as an unopposed CaN activity can dephosphorylate LTCCs ^14^. This in addition to the presence of fewer CP-AMPARs in LHb neurons of ΔPKA mice might result in further reduction in calcium influx, defective eCB production, and hence impaired eCB-LTD in LHb neurons.

GABA_A_Rs at GABAergic synapses onto LHb neurons are mainly composed of a combination of the α1-3, β1 and γ1-2 subunits^75^. There is less known about PKA-dependent regulation of GABA_A_Rs in the LHb. Our previous study in VTA DA neurons suggests that activation of dopamine D2 receptors results in PKA inhibition that promotes AKAP150-CaN-mediated internalization of GABA_A_R receptors and the expression of LTD at GABAergic synapses onto VTA DA neurons^29^. The expression of an inhibitory mGluR-dependent postsynaptic LTD at GABAergic synapses onto LHb neurons requires a PKC-dependent phosphorylation of the β2 receptor subunits of GABA_A_Rs, reducing GABA_A_R single-channel conductance^60^. This also excludes the possibility that a biased AKAP150-PKC signaling in the absence of AKAP-PKA association in ΔPKA mice could promote the basal increase in the conductance of GABA_A_Rs in LHb neurons. Therefore, it is still an open question which AKAP150 associations with other binding partners could promote forward trafficking of GABA_A_Rs in LHb neurons.

In addition to alterations of synaptic transmission and LTD by ΔPKA mutation, we observed a significant increase in LHb intrinsic excitability associated with higher input resistance and lower amplitude of mAHPs, mimicking the effects of exogenous CRF (as we observed in both mouse and rat LHb) and after a severe early like stress (i.e., maternal deprivation)^58^. However, the diminishing effects of exogenous CRF and maternal deprivation on mAHPs and the resultant hyperexcitability were due to the PKA-dependent decrease in the function and/or abundance of SK channels^58^, a mechanism that is less likely to underlie ΔPKA mutation-induced LHb intrinsic plasticity. Afterhyperopolarizations including fAHPs and mAHPs are mediated by diverse types of potassium channels that repolarize the membrane to regulate and limit excessive neuronal excitability. In addition to SK channels, voltage gated K^+^ channel 7 (Kv7, also known as M currents) contribute to mAHP in neurons^76^. Therefore, it is possible that genetic disruption of AKAP150 anchoring of PKA in ΔPKA favors AKAP150-anchored PKC and the resultant inhibition of M-type mAHPs ^12^ to increase LHb intrinsic excitability in ΔPKA mice, which could saturate and occlude the excitatory actions of CRF on LHb intrinsic excitability. Consistent with this interpretation, it has been shown that activation of LHb M channels reduces LHb neuronal activity and blocks the anxiety-like phenotype in alcohol-withdrawn rats^77^. Given that that the majority of synaptic inputs to the LHb co-release glutamate and GABA^41^, our observation of CRF-induced suppression of both presynaptic glutamate and GABA release in mouse LHb is not surprising as CB1R are expressed on presynaptic terminals in the LHb where CB1R activation by eCBs can reduce the probability of presynaptic glutamate and GABA release onto LHb neurons at distinct synaptic inputs to the LHb (e.g., LPO) although the effect on presynaptic GABA release is assumed to be predominantly larger ^78^. ΔPKA mutation to some extent reduced the suppressing effects of CRF on presynaptic GABA release but also unmasked a small potentiating effect of CRF on presynaptic glutamate release. This could be due to decreased depolarization and/ or calcium influx from the fewer CP-AMPARs available at the synapse as well as the less effective influx of calcium from hypofunctional LTCC in ΔPKA mice, which could in turn lead to dysregulation of eCB production that not only prevented the expression of eCB-LTD but also blunted the inhibitory effects of CRF on synaptic transmission by shifting excitation/inhibition balance to more excitation. Therefore, we assume that disruption of AKAP150 anchoring of PKA seems to promote LHb hyperexcitability through synaptic and intrinsic mechanisms that may relate to the lack of AKAP150-dependent PKA-mediated signaling as well as favoring unopposed non-PKA-mediated AKAP150 interactions. These concepts are briefly summarized in Fig 11, which depicts in schematic form representative GABAergic and glutamatergic terminals innervating a spine and dendritic shaft of an LHb neuron. Sites in this synaptic complex where AKAP-mediated signaling plays a potential role in synaptic function and plasticity are indicated.

**Figure 11.**
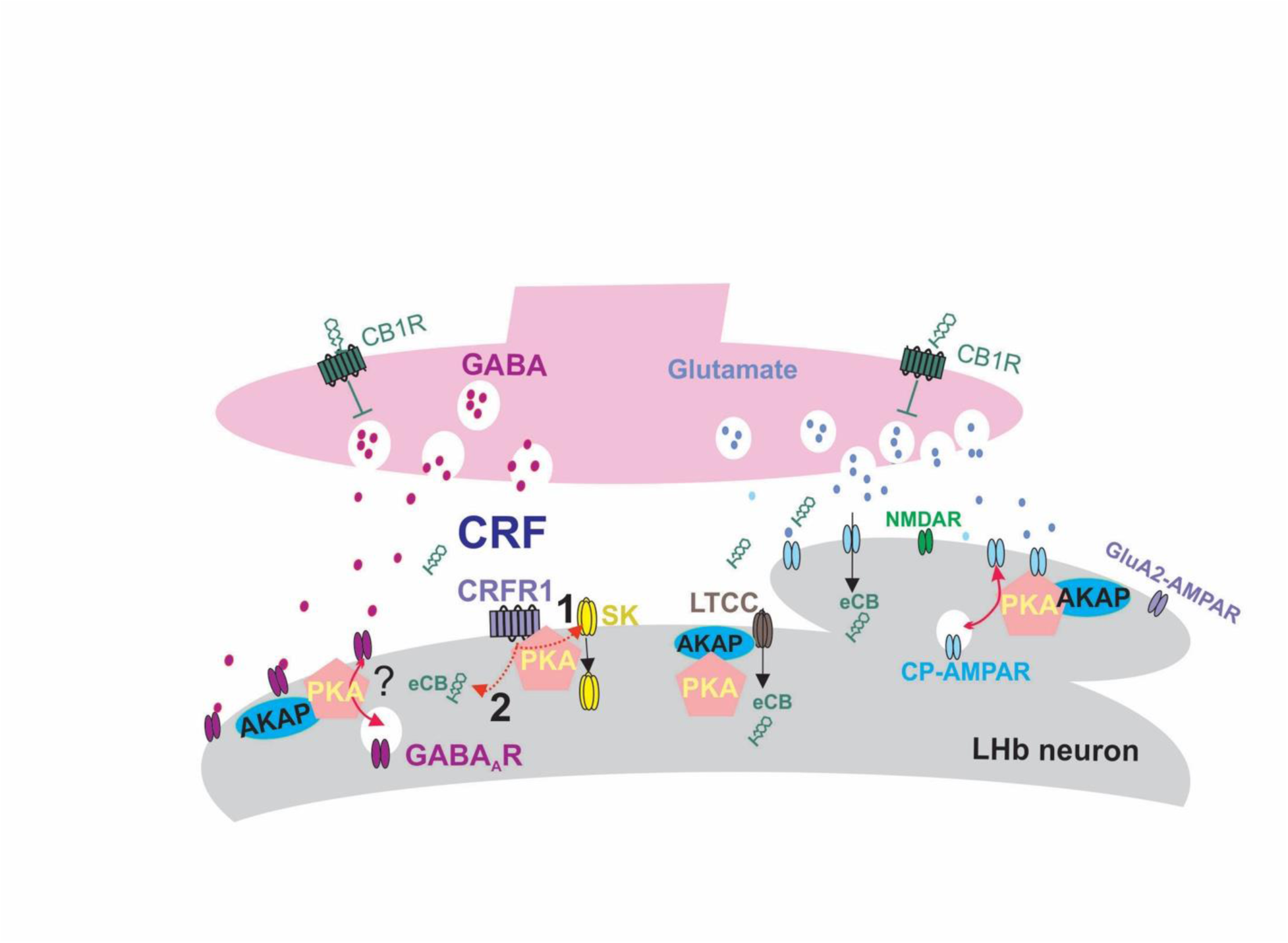
Schematic proposed model illustrating regulatory roles of AKAP150-anchored PKA signaling in synaptic function, LTD and intrinsic excitability of LHb neurons. This model predicts that AKAP150-PKA association is necessary for maintenance of AMPARs at glutamatergic synapses and expression of LTD by LFS. We assume that the sources of increased depolarization and calcium influx for eCB production during LTD induction may arise from CP AMPARs and/or LTCC both regulated by AKAP150-PKA complex. Therefore, disruption of AKAP150 anchoring of PKA to CP AMPARs in LHb neurons in ΔPKA mice results in lower number of CP AMPARs at glutamatergic synapses. The reduced influx of calcium from fewer available CP AMPARs at the synapse as well as hypofunctional LTCC by the genetic disruption of PKA-AKAP150 association can impair eCB production and thus the expression of eCB-LTD. Our model also predicts that defective AKAP150-PKA association may reduce trafficking and/or function of potassium channels mediating mAHPs (e.g., M currents, not shown). The model also shows the known effects of CRF-CRFR1-PKA signaling in the LHb which results in eCB production as well as promote LHb hyperexcitability through modulation of trafficking or conductance of SK potassium channels. Thus, we assume that under pathological conditions AKAP150-PKA dysregulation of CP AMPARs, LTCC, M currents and eCB signaling could promote LHb hyperexcitability and blunt CRF neuromodulatory actions. While our model may indicate the existence of inhibitory effects of AKAP150-PKA on GABA_A_R feedforward trafficking, it is still unclear which AKAP150 partners may contribute to regulation of GABA_A_Rs in LHb neurons. Theindicates such unresolved questions.

Overall, our study highlights significant and multifaceted impacts of AKAP150 anchoring of PKA in regulation of glutamatergic transmission and plasticity, and neuronal excitability of LHb neurons. Moreover, defects in AKAP150-mediated PKA anchoring under pathological processes could favor other, yet to be discovered, AKAP150 interactions in LHb neurons that promote LHb hyperactivity and dysregulate CRF neuromodulation within the LHb, reminiscent of the effects of a severe early life stress^58^ and alcohol withdrawal^79, 80^. Future studies are needed to increase our understanding of cell-type and input-specific neuroplasticity and neuromodulation of LHb activity through alterations in LHb AKAP150 complex interactions in neurological and neuropsychiatric illnesses.

## Acknowledgements

The opinions and assertions contained herein are the private opinions of the authors and are not to be construed as official or reflecting the views of the Uniformed Services University of the Health Sciences or the Department of Defense or the Government of the United States.

## Author contributions

FSN and MLD designed the research; SCS, WJF, LDL, RDS and CB performed electrophysiology; JLS was responsible for breeding ΔPKA mice; SG KMG and EHT performed immunohistochemistry. SCS, WJF and FSN analyzed the data and prepared the figures; SCS, WJF, BMC, MLD and FSN wrote the initial draft of the manuscript. All authors critically reviewed the content and approved final version of manuscript for submission.

## Data Sharing

The data that support the findings of this study are available on request from the corresponding authors. The data are not publicly available due to privacy or ethical restrictions.

## References

1. Sanderson, J.L. and Dell’acqua, M.L. (2011). AKAP Signaling Complexes in Regulation of Excitatory Synaptic Plasticity. Neuroscientist 17, 321–336.

2. Snyder, E.M., Colledge, M., Crozier, R.A., Chen, W.S., Scott, J.D. and Bear, M.F. (2005). Role for A kinase-anchoring proteins (AKAPS) in glutamate receptor trafficking and long term synaptic depression. J Biol Chem 280, 16962–16968.

3. Lu, Y., Allen, M., Halt, A.R., Weisenhaus, M., Dallapiazza, R.F., Hall, D.D., Usachev, Y.M., McKnight, G.S. and Hell, J.W. (2007). Age-dependent requirement of AKAP150-anchored PKA and GluR2-lacking AMPA receptors in LTP. EMBO J 26, 4879–4890.

4. Lu, Y., Zhang, M., Lim, I.A., Hall, D.D., Allen, M., Medvedeva, Y., McKnight, G.S., Usachev, Y.M. and Hell, J.W. (2008). AKAP150-anchored PKA activity is important for LTD during its induction phase. The Journal of physiology 586, 4155–4164.

5. Bhattacharyya, S., Biou, V., Xu, W., Schluter, O. and Malenka, R.C. (2009). A critical role for PSD-95/AKAP interactions in endocytosis of synaptic AMPA receptors. Nature neuroscience 12, 172–181.

6. Jurado, S., Biou, V. and Malenka, R.C. (2011). A calcineurin/AKAP complex is required for NMDA receptor-dependent long-term depression. Nature neuroscience 13, 1053–1055.

7. Sanderson, J.L., Gorski, J.A., Gibson, E.S., Lam, P., Freund, R.K., Chick, W.S. and Dell’Acqua, M.L. (2012). AKAP150-anchored calcineurin regulates synaptic plasticity by limiting synaptic incorporation of Ca2+-permeable AMPA receptors. J Neurosci 32, 15036–15052.

8. Authement, M.E., Kodangattil, J.N., Gouty, S., Rusnak, M., Symes, A.J., Cox, B.M. and Nugent, F.S. (2015). Histone Deacetylase Inhibition Rescues Maternal Deprivation-Induced GABAergic Metaplasticity through Restoration of AKAP Signaling. Neuron 86, 1240–1252.

9. Sanderson, J.L., Gorski, J.A. and Dell’Acqua, M.L. (2016). NMDA Receptor-Dependent LTD Requires Transient Synaptic Incorporation of Ca(2)(+)-Permeable AMPARs Mediated by AKAP150-Anchored PKA and Calcineurin. Neuron 89, 1000–1015.

10. Zhang, J. and Shapiro, M.S. (2012). Activity-dependent transcriptional regulation of M-Type (Kv7) K(+) channels by AKAP79/150-mediated NFAT actions. Neuron 76, 1133–1146.

11. Hoshi, N., Langeberg, L.K. and Scott, J.D. (2005). Distinct enzyme combinations in AKAP signalling complexes permit functional diversity. Nat Cell Biol 7, 1066–1073.

12. Hoshi, N., Zhang, J.S., Omaki, M., Takeuchi, T., Yokoyama, S., Wanaverbecq, N., Langeberg, L.K., Yoneda, Y., Scott, J.D., Brown, D.A. and Higashida, H. (2003). AKAP150 signaling complex promotes suppression of the M-current by muscarinic agonists. Nature neuroscience 6, 564–571.

13. Lin, L., Sun, W., Kung, F., Dell’Acqua, M.L. and Hoffman, D.A. (2011). AKAP79/150 impacts intrinsic excitability of hippocampal neurons through phospho-regulation of A-type K+ channel trafficking. J Neurosci 31, 1323–1332.

14. Murphy, J.G., Sanderson, J.L., Gorski, J.A., Scott, J.D., Catterall, W.A., Sather, W.A. and Dell’Acqua, M.L. (2014). AKAP-Anchored PKA Maintains Neuronal L-type Calcium Channel Activity and NFAT Transcriptional Signaling. Cell reports.

15 Bai, X., Zhang, K., Ou, C., Nie, B., Zhang, J., Huang, Y., Zhang, Y., Huang, J., Ouyang, H., Cao, M. and Huang, W. (2023). Selective activation of AKAP150/TRPV1 in ventrolateral periaqueductal gray GABAergic neurons facilitates conditioned place aversion in male mice. Commun Biol 6, 742.

16. Schnizler, K., Shutov, L.P., Van Kanegan, M.J., Merrill, M.A., Nichols, B., McKnight, G.S., Strack, S., Hell, J.W. and Usachev, Y.M. (2008). Protein kinase A anchoring via AKAP150 is essential for TRPV1 modulation by forskolin and prostaglandin E2 in mouse sensory neurons. J Neurosci 28, 4904–4917.

17. Tavalin, S.J., Colledge, M., Hell, J.W., Langeberg, L.K., Huganir, R.L. and Scott, J.D. (2002). Regulation of GluR1 by the A-kinase anchoring protein 79 (AKAP79) signaling complex shares properties with long-term depression. J Neurosci 22, 3044–3051.

18. Purkey, A.M. and Dell’Acqua, M.L. (2020). Phosphorylation-Dependent Regulation of Ca(2+)-Permeable AMPA Receptors During Hippocampal Synaptic Plasticity. Front Synaptic Neurosci 12, 8.

19. Sanderson, J.L., Scott, J.D. and Dell’Acqua, M.L. (2018). Control of Homeostatic Synaptic Plasticity by AKAP-Anchored Kinase and Phosphatase Regulation of Ca(2+)-Permeable AMPA Receptors. J Neurosci 38, 2863–2876.

20. Richter, S., Gorny, X., Marco-Pallares, J., Kramer, U.M., Machts, J., Barman, A., Bernstein, H.G., Schule, R., Schols, L., Rodriguez-Fornells, A., Reissner, C., Wustenberg, T., Heinze, H.J., Gundelfinger, E.D., Duzel, E., Munte, T.F., Seidenbecher, C.I. and Schott, B.H. (2011). A Potential Role for a Genetic Variation of AKAP5 in Human Aggression and Anger Control. Frontiers in human neuroscience 5, 175.

21. Suryavanshi, S.V., Jadhav, S.M. and McConnell, B.K. (2018). Polymorphisms/Mutations in A-Kinase Anchoring Proteins (AKAPs): Role in the Cardiovascular System. J Cardiovasc Dev Dis 5.

22. Richter, S., Gorny, X., Machts, J., Behnisch, G., Wustenberg, T., Herbort, M.C., Munte, T.F., Seidenbecher, C.I. and Schott, B.H. (2013). Effects of AKAP5 Pro100Leu genotype on working memory for emotional stimuli. PLoS One 8, e55613.

23. Sutrala, S.R., Goossens, D., Williams, N.M., Heyrman, L., Adolfsson, R., Norton, N., Buckland, P.R. and Del-Favero, J. (2007). Gene copy number variation in schizophrenia. Schizophr Res 96, 93–99.

24. Scheller-Gilkey, G., Moynes, K., Cooper, I., Kant, C. and Miller, A.H. (2004). Early life stress and PTSD symptoms in patients with comorbid schizophrenia and substance abuse. Schizophr Res 69, 167–174.

25. Scheller-Gilkey, G., Thomas, S.M., Woolwine, B.J. and Miller, A.H. (2002). Increased early life stress and depressive symptoms in patients with comorbid substance abuse and schizophrenia. Schizophrenia bulletin 28, 223–231.

26. Winklbaur, B., Ebner, N., Sachs, G., Thau, K. and Fischer, G. (2006). Substance abuse in patients with schizophrenia. Dialogues in clinical neuroscience 8, 37–43.

27. Shepard, R.D., Langlois, L.D., Authement, M.E. and Nugent, F.S. (2020). Histone deacetylase inhibition reduces ventral tegmental area dopamine neuronal hyperexcitability involving AKAP150 signaling following maternal deprivation in juvenile male rats. J Neurosci Res 98, 1457–1467.

28. Shepard, R.D., Gouty, S., Kassis, H., Berenji, A., Zhu, W., Cox, B.M. and Nugent, F.S. (2018). Targeting histone deacetylation for recovery of maternal deprivation-induced changes in BDNF and AKAP150 expression in the VTA. Experimental neurology 309, 160–168.

29. Dacher, M., Gouty, S., Dash, S., Cox, B.M. and Nugent, F.S. (2013). A-kinase anchoring protein-calcineurin signaling in long-term depression of GABAergic synapses. J Neurosci 33, 2650–2660.

30. Reissner, K.J., Uys, J.D., Schwacke, J.H., Comte-Walters, S., Rutherford-Bethard, J.L., Dunn, T.E., Blumer, J.B., Schey, K.L. and Kalivas, P.W. (2011). AKAP Signaling in Reinstated Cocaine Seeking Revealed by iTRAQ Proteomic Analysis. J Neurosci 31, 5648–5658.

31. Guercio, L.A., Hofmann, M.E., Swinford-Jackson, S.E., Sigman, J.S., Wimmer, M.E., Dell’Acqua, M.L., Schmidt, H.D. and Pierce, R.C. (2018). A-Kinase Anchoring Protein 150 (AKAP150) Promotes Cocaine Reinstatement by Increasing AMPA Receptor Transmission in the Accumbens Shell. Neuropsychopharmacology 43, 1395–1404.

32. Bai, X., Zhang, K., Ou, C., Mu, Y., Chi, D., Zhang, J., Huang, J., Li, X., Zhang, Y., Huang, W. and Ouyang, H. (2023). AKAP150 from nucleus accumbens dopamine D1 and D2 receptor-expressing medium spiny neurons regulates morphine withdrawal. iScience 26, 108227.

33. Zhou, H.Y., He, J.G., Hu, Z.L., Xue, S.G., Xu, J.F., Cui, Q.Q., Gao, S.Q., Zhou, B., Wu, P.F., Long, L.H., Wang, F. and Chen, J.G. (2019). A-Kinase Anchoring Protein 150 and Protein Kinase A Complex in the Basolateral Amygdala Contributes to Depressive-like Behaviors Induced by Chronic Restraint Stress. Biological psychiatry 86, 131–142.

34. Moita, M.A., Lamprecht, R., Nader, K. and LeDoux, J.E. (2002). A-kinase anchoring proteins in amygdala are involved in auditory fear memory. Nature neuroscience 5, 837–838.

35. Hu, H., Cui, Y. and Yang, Y. (2020). Circuits and functions of the lateral habenula in health and in disease. Nat Rev Neurosci 21, 277–295.

36. Proulx, C.D., Hikosaka, O. and Malinow, R. (2014). Reward processing by the lateral habenula in normal and depressive behaviors. Nature neuroscience 17, 1146–1152.

37. Hikosaka, O. (2010). The habenula: from stress evasion to value-based decision-making. Nat Rev Neurosci 11, 503–513.

38. Baker, P.M., Mathis, V., Lecourtier, L., Simmons, S.C., Nugent, F.S., Hill, S. and Mizumori, S.J.Y. (2022). Lateral Habenula Beyond Avoidance: Roles in Stress, Memory, and Decision-Making With Implications for Psychiatric Disorders. Front Syst Neurosci 16, 826475.

39. Proulx, C.D., Hikosaka, O. and Malinow, R. (2014). Reward processing by the lateral habenula in normal and depressive behaviors. Nature neuroscience 17, 1146–1152.

40. Zhang, L., Hernandez, V.S., Swinny, J.D., Verma, A.K., Giesecke, T., Emery, A.C., Mutig, K., Garcia-Segura, L.M. and Eiden, L.E. (2018). A GABAergic cell type in the lateral habenula links hypothalamic homeostatic and midbrain motivation circuits with sex steroid signaling. Translational psychiatry 8, 50.

41. Webster, J.F., Vroman, R., Balueva, K., Wulff, P., Sakata, S. and Wozny, C. (2020). Disentangling neuronal inhibition and inhibitory pathways in the lateral habenula. Sci Rep 10, 8490.

42. Zhang, L., Hernandez, V.S., Vazquez-Juarez, E., Chay, F.K. and Barrio, R.A. (2016). Thirst Is Associated with Suppression of Habenula Output and Active Stress Coping: Is there a Role for a Non-canonical Vasopressin-Glutamate Pathway? Frontiers in neural circuits 10, 13.

43. Margolis, E.B. and Fields, H.L. (2016). Mu Opioid Receptor Actions in the Lateral Habenula. PLoS One 11, e0159097.

44. Graziane, N.M., Neumann, P.A. and Dong, Y. (2018). A Focus on Reward Prediction and the Lateral Habenula: Functional Alterations and the Behavioral Outcomes Induced by Drugs of Abuse. Front Synaptic Neurosci 10, 12.

45. Stopper, C.M., Tse, M.T.L., Montes, D.R., Wiedman, C.R. and Floresco, S.B. (2014). Overriding phasic dopamine signals redirects action selection during risk/reward decision making. Neuron 84, 177–189.

46. Quina, L.A., Tempest, L., Ng, L., Harris, J.A., Ferguson, S., Jhou, T.C. and Turner, E.E. (2015). Efferent Pathways of the Mouse Lateral Habenula. Journal of Comparative Neurology 523, 32–60.

47. Proulx, C.D., Aronson, S., Milivojevic, D., Molina, C., Loi, A., Monk, B., Shabel, S.J. and Malinow, R. (2018). A neural pathway controlling motivation to exert effort. Proceedings of the National Academy of Sciences of the United States of America 115, 5792–5797.

48. van Zessen, R., Phillips, J.L., Budygin, E.A. and Stuber, G.D. (2012). Activation of VTA GABA Neurons Disrupts Reward Consumption. Neuron 73, 1184–1194.

49. Cerniauskas, I., Winterer, J., de Jong, J.W., Lukacsovich, D., Yang, H., Khan, F., Peck, J.R., Obayashi, S.K., Lilascharoen, V., Lim, B.K., Foldy, C. and Lammel, S. (2019). Chronic Stress Induces Activity, Synaptic, and Transcriptional Remodeling of the Lateral Habenula Associated with Deficits in Motivated Behaviors. Neuron 104, 899–915 e898.

50. Pobbe, R.L. and Zangrossi, H., Jr. (2008). Involvement of the lateral habenula in the regulation of generalized anxiety- and panic-related defensive responses in rats. Life sciences 82, 1256–1261.

51. Berger, A.L., Henricks, A.M., Lugo, J.M., Wright, H.R., Warrick, C.R., Sticht, M.A., Morena, M., Bonilla, I., Laredo, S.A., Craft, R.M., Parsons, L.H., Grandes, P.R., Hillard, C.J., Hill, M.N. and McLaughlin, R.J. (2018). The Lateral Habenula Directs Coping Styles Under Conditions of Stress via Recruitment of the Endocannabinoid System. Biological psychiatry 84, 611–623.

52. Lee, Y.-A. and Goto, Y. (2021). The Habenula in the Link Between ADHD and Mood Disorder. Frontiers in Behavioral Neuroscience 15.

53. Li, J., Yang, S., Liu, X., Han, Y., Li, Y., Feng, J. and Zhao, H. (2019). Hypoactivity of the lateral habenula contributes to negative symptoms and cognitive dysfunction of schizophrenia in rats. Experimental neurology 318, 165–173.

54. Arfuso, M., Salas, R., Castellanos, F.X. and Krain Roy, A. (2021). Evidence of Altered Habenular Intrinsic Functional Connectivity in Pediatric ADHD. J Atten Disord 25, 749–757.

55. Orsini, C.A., Moorman, D.E., Young, J.W., Setlow, B. and Floresco, S.B. (2015). Neural mechanisms regulating different forms of risk-related decision-making: Insights from animal models. Neuroscience and biobehavioral reviews 58, 147–167.

56. Banwinkler, M., Theis, H., Prange, S. and van Eimeren, T. (2022). Imaging the Limbic System in Parkinson’s Disease-A Review of Limbic Pathology and Clinical Symptoms. Brain Sci 12.

57. Zhu, Y., Qi, S., Zhang, B., He, D., Teng, Y., Hu, J. and Wei, X. (2019). Connectome-Based Biomarkers Predict Subclinical Depression and Identify Abnormal Brain Connections With the Lateral Habenula and Thalamus. Frontiers in psychiatry 10.

58. Authement, M.E., Langlois, L.D., Shepard, R.D., Browne, C.A., Lucki, I., Kassis, H. and Nugent, F.S. (2018). A role for corticotropin-releasing factor signaling in the lateral habenula and its modulation by early-life stress. Science signaling 11.

59. Flerlage, W.J., Langlois, L.D., Rusnak, M., Simmons, S.C., Gouty, S., Armstrong, R.C., Cox, B.M., Symes, A.J., Tsuda, M.C. and Nugent, F.S. (2023). Involvement of Lateral Habenula Dysfunction in Repetitive Mild Traumatic Brain Injury-Induced Motivational Deficits. J Neurotrauma 40, 125–140.

60. Valentinova, K. and Mameli, M. (2016). mGluR-LTD at Excitatory and Inhibitory Synapses in the Lateral Habenula Tunes Neuronal Output. Cell reports 16, 2298–2307.

61. Park, H., Rhee, J., Lee, S. and Chung, C. (2017). Selectively Impaired Endocannabinoid-Dependent Long-Term Depression in the Lateral Habenula in an Animal Model of Depression. Cell reports 20, 289–296.

62. Kang, M., Noh, J. and Chung, J.M. (2020). NMDA receptor-dependent long-term depression in the lateral habenula: implications in physiology and depression. Sci Rep 10, 17921.

63. Wild, A.R. and Dell’Acqua, M.L. (2017). Potential for therapeutic targeting of AKAP signaling complexes in nervous system disorders. Pharmacol Ther.

64. Herring, D., Huang, R., Singh, M., Dillon, G.H. and Leidenheimer, N.J. (2005). PKC modulation of GABAA receptor endocytosis and function is inhibited by mutation of a dileucine motif within the receptor beta 2 subunit. Neuropharmacology 48, 181–194.

65. Gutknecht, E., Vauquelin, G. and Dautzenberg, F.M. (2010). Corticotropin-releasing factor receptors induce calcium mobilization through cross-talk with Gq-coupled receptors. European journal of pharmacology 642, 1–9.

66. Dautzenberg, F.M., Gutknecht, E., Van der Linden, I., Olivares-Reyes, J.A., Dürrenberger, F. and Hauger, R.L. (2004). Cell-type specific calcium signaling by corticotropin-releasing factor type 1 (CRF1) and 2a (CRF2(a)) receptors: phospholipase C-mediated responses in human embryonic kidney 293 but not SK-N-MC neuroblastoma cells. Biochem Pharmacol 68, 1833–1844.

67. West, A.E. and Greenberg, M.E. (2011). Neuronal activity-regulated gene transcription in synapse development and cognitive function. Cold Spring Harb Perspect Biol 3.

68. Maroteaux, M. and Mameli, M. (2012). Cocaine evokes projection-specific synaptic plasticity of lateral habenula neurons. J Neurosci 32, 12641–12646.

69. Li, B., Piriz, J., Mirrione, M., Chung, C., Proulx, C.D., Schulz, D., Henn, F. and Malinow, R. (2011). Synaptic potentiation onto habenula neurons in the learned helplessness model of depression. Nature 470, 535–539.

70. Bucko, P.J. and Scott, J.D. (2021). Drugs That Regulate Local Cell Signaling: AKAP Targeting as a Therapeutic Option. Annual review of pharmacology and toxicology 61, 361–379.

71. Meye, F.J., Valentinova, K., Lecca, S., Marion-Poll, L., Maroteaux, M.J., Musardo, S., Moutkine, I., Gardoni, F., Huganir, R.L., Georges, F. and Mameli, M. (2015). Cocaine-evoked negative symptoms require AMPA receptor trafficking in the lateral habenula. Nature neuroscience 18, 376–378.

72. Li, K., Zhou, T., Liao, L., Yang, Z., Wong, C., Henn, F., Malinow, R., Yates, J.R., 3rd and Hu, H. (2013). betaCaMKII in lateral habenula mediates core symptoms of depression. Science (New York, N.Y 341, 1016–1020.

73. Kerchner, G.A. and Nicoll, R.A. (2008). Silent synapses and the emergence of a postsynaptic mechanism for LTP. Nat Rev Neurosci 9, 813–825.

74. Woolfrey, K.M., O’Leary, H., Goodell, D.J., Robertson, H.R., Horne, E.A., Coultrap, S.J., Dell’Acqua, M.L. and Bayer, K.U. (2018). CaMKII regulates the depalmitoylation and synaptic removal of the scaffold protein AKAP79/150 to mediate structural long-term depression. J Biol Chem 293, 1551–1567.

75. Hörtnagl, H., Tasan, R.O., Wieselthaler, A., Kirchmair, E., Sieghart, W. and Sperk, G. (2013). Patterns of mRNA and protein expression for 12 GABAA receptor subunits in the mouse brain. Neuroscience 236, 345–372.

76. Dwivedi, D. and Bhalla, U.S. (2021). Physiology and Therapeutic Potential of SK, H, and M Medium AfterHyperPolarization Ion Channels. Frontiers in molecular neuroscience 14, 658435.

77. Kang, S., Li, J., Zuo, W., Fu, R., Gregor, D., Krnjevic, K., Bekker, A. and Ye, J.-H. (2017). Ethanol Withdrawal Drives Anxiety-Related Behaviors by Reducing M-type Potassium Channel Activity in the Lateral Habenula. Neuropsychopharmacology 42, 1813–1824.

78. Winters, N.D., Kondev, V., Loomba, N., Delpire, E., Grueter, B.A. and Patel, S. (2023). Opposing retrograde and astrocyte-dependent endocannabinoid signaling mechanisms regulate lateral habenula synaptic transmission. Cell reports 42, 112159.

79. Zuo, W., Zuo, Q., Wu, L., Mei, Q., Shah, M., Zheng, J., Li, D., Xu, Y. and Ye, J.H. (2021). Roles of corticotropin-releasing factor signaling in the lateral habenula in anxiety-like and alcohol drinking behaviors in male rats. Neurobiol Stress 15, 100395.

80. Fu, R., Mei, Q., Shiwalkar, N., Zuo, W., Zhang, H., Gregor, D., Patel, S. and Ye, J.H. (2020). Anxiety during alcohol withdrawal involves 5-HT2C receptors and M-channels in the lateral habenula. Neuropharmacology 163, 107863.

